# Identification of RNA Targets of Classical and Non-Canonical RNA-binding Proteins by soniCLIP

**DOI:** 10.64898/2026.08.17.745202

**Authors:** Pia Sommerkamp, Sudeep Sahadevan, Thileepan Sekaran, Silvia Colucci, Dunja Ferring-Appel, Matthias W. Hentze

## Abstract

GRAPHICAL ABSTRACT

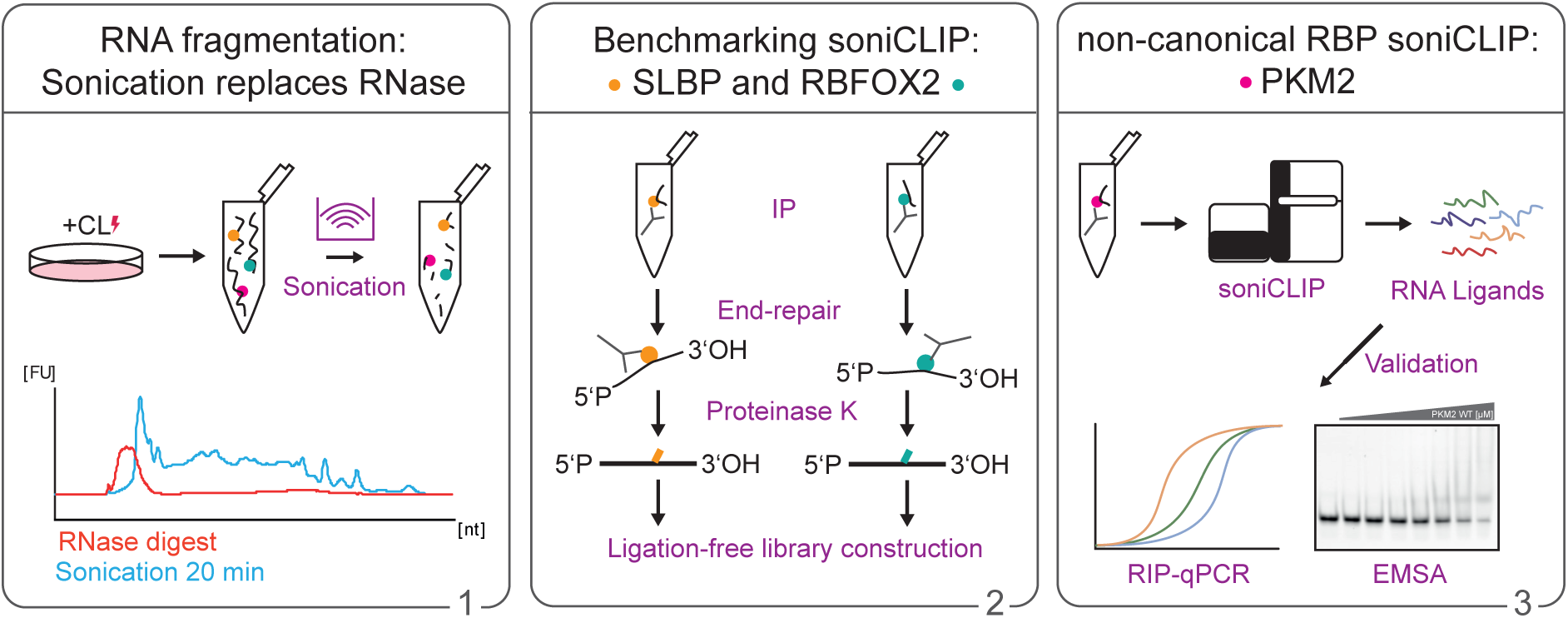

Crosslinking and immunoprecipitation followed by sequencing (CLIP-seq) is widely used to identify the RNA targets of RNA-binding proteins (RBPs). However, its application to non-canonical RBPs lacking canonical RNA-binding domains and frequently displaying low or transient RNA occupancy, is limited by low signal-to-noise ratios, high input requirements and error-prone ligation steps during library preparation. To overcome these limitations, we developed soniCLIP, a streamlined CLIP-seq workflow that replaces RNase-mediated RNA fragmentation with sonication and uses a ligation-free strategy for library construction. soniCLIP is optimized for reproducible identification of enriched RBP-associated RNA regions from limited starting material. We benchmarked soniCLIP against the widespread eCLIP approach and observed reproducible recovery of known RBP-associated regions and target recovery comparable to ENCODE eCLIP, while requiring only 10% (500 µg) of protein input. We further applied soniCLIP to the glycolytic enzyme and non-canonical RBP pyruvate kinase M2 (PKM2). We identified 197 significantly enriched RNA regions and validated selected targets by RIP-qRT-PCR and *in vitro* binding assays. By combining reduced input requirements, high reproducibility, a shortened 3.5-day workflow and the elimination of gel-based purification, soniCLIP provides an efficient and robust approach for the identification of RNA targets of canonical and non-canonical RBPs.

## INTRODUCTION

CLIP-seq has become a key strategy to define RNA targets and binding sites of RNA-binding proteins (RBPs) in cells. CLIP relies on ultraviolet (UV)-mediated covalent stabilization of protein-RNA contacts, followed by RNA fragmentation (typically by limited RNase digestion), immunoprecipitation (IP) of an RBP of interest, recovery of associated RNA fragments, sequencing library preparation and computational identification of enriched RNA regions (1–3). This protein-centric strategy has enabled transcriptome-wide analyses of RNA-protein interactions in their cellular context and has become central to the functional annotation of RBPs. Since the first CLIP experiments (1), numerous variants have been developed, including HITS-CLIP, PAR-CLIP, iCLIP and enhanced CLIP (eCLIP), which differ in crosslink detection, RNA processing, library construction, background correction and binding-site resolution (4–7) (reviewed in (2)).

Despite these advances, CLIP remains experimentally and computationally demanding, with several critical sources of material loss and bias arising after crosslinking (reviewed in (2)). RNA fragmentation is particularly consequential and directly affects read mapping and binding-site assignment (8). RNase digestion must therefore be carefully optimized: overdigestion can generate fragments that are too short to map reliably, whereas underdigestion can yield fragments that are too long for efficient library construction and precise localization of binding regions (9,10). Moreover, we show that RNase treatment can influence IP specificity, particularly for non-canonical RBPs, by contributing to the co-recovery of nonspecific RNA fragments. Subsequent gel- or membrane-based purification of RBP-RNA complexes increases hands-on time and can limit recovery, accessibility and scalability (11,12). Finally, ligation-dependent library preparation introduces additional sources for sample loss and adapter-related artifacts, including ligation of fragments or double-ligation events, which reduce the number of informative reads available for downstream data analysis (4). This loss is not merely a computational inconvenience; it reflects material attrition and technical bias that, across available CLIP technologies, leaves only 13-57% of input reads aligned to the genome for statistical analysis of the binding sites (2). Thus, although CLIP-based approaches have transformed the analysis of RBP target landscapes, their performance remains constrained by nonspecific RNA fragments, labor-intensive purification steps and losses introduced during library construction.

These constraints are particularly consequential when starting material is limited or when only a small fraction of the protein under study is RNA-bound. Lower input requirements would broaden CLIP applications to scarce cell populations, primary material, sorted cells, organoids and perturbation settings in which sample amounts are small. Low RNA occupancy is also expected for many non-canonical RBPs, which lack high affinity, globular RNA-binding domains (RBDs) (13–19). Their RNA binding may be transient, condition-dependent or limited to a minor fraction of the cellular protein pool, and many do not display clear sequence-specific binding preferences (19). For such proteins, limited recovery of RNA-protein material increases the impact of background and makes CLIP analyses especially sensitive to losses introduced during IP and library construction.

We therefore developed soniCLIP, a streamlined CLIP-seq workflow for identifying RNA targets of canonical and non-canonical RBPs. soniCLIP replaces RNase-mediated fragmentation with sonication, generating RNA fragments in a sequencing-compatible size range while eliminating RNase-associated background in IP samples. It further uses ligation-free library construction to avoid adapter-ligation artifacts and associated read loss. Together, these changes simplify the workflow, eliminate gel-based purification, shorten the protocol to 3.5 days (from 5 days for standard eCLIP and iCLIP protocols) and reduce the required input material to as little as 500 µg of protein starting material.

To benchmark soniCLIP against the widely-used eCLIP protocol (4,20), we first selected two well-characterized canonical RBPs with established binding preferences and extensive public CLIP resources: Stem-Loop Binding Protein (SLBP) and RNA-binding Fox-1 homolog 2 (RBFOX2). SLBP binds the conserved stem-loop element at the 3′ end of replication-dependent histone mRNAs (21), whereas RBFOX2 recognizes defined RNA sequence motifs (22). We then applied soniCLIP to pyruvate kinase M2 (PKM2), a glycolytic enzyme and representative non-canonical RBP identified in several interactome studies (15,16,18,23). PKM2 has been linked to RNA-associated processes, including RNA G-quadruplex binding, ribosome association and regulation of translation and mRNA fate (24–26). As an abundant metabolic enzyme for which only a small fraction of the cellular protein pool may be RNA-bound, PKM2 represents a biologically relevant and technically challenging model for evaluating CLIP performance on non-canonical RBPs.

Here, we show that soniCLIP reproducibly identifies established target regions of canonical RBPs at least as well as eCLIP, and enables RNA target discovery for the non-canonical RBP PKM2 from low-input material. By reducing protocol complexity, avoiding ligation-associated read loss and improving accessibility, soniCLIP provides a robust approach for mapping RNA targets of RBPs across a broad range of RNA-binding modes.

## MATERIALS AND METHODS

### MATERIALS

Antibodies

| Antibody | Supplier | Clone/Catalog # |
| --- | --- | --- |
| ACTIN | Sigma-Aldrich | Monoclonal AC-15; #A5441 |
| PKM2 | Cell Signaling | Monoclonal D78A4; #13266S |
| HuR | Proteintech | Polyclonal; #11910-1-AP |
| Histone H3 | Proteintech | Polyclonal; #17168-1-AP |
| SLBP | Thermo Fisher | Polyclonal; #A303-968A |
| RBFOX2 | Thermo Fisher | Polyclonal; #A300-864A |
| Rabbit IgG control | Invitrogen | Polyclonal; #10500C |
| HRP-conjugated anti-mouse IgG | Abcam | Polyclonal; #AB6789 |
| HRP-conjugated anti-rabbit IgG | Abcam | Polyclonal; #AB97051 |
| HRP-conjugated anti-rabbit IgG, light chain specific | Proteintech | Monoclonal; #SA00001-7L |

Enzymes

| Enzyme | Supplier | Catalog # |
| --- | --- | --- |
| Turbo DNase | Thermo Fisher/Invitrogen | #AM2238 |
| Ambion RNase I | Thermo Fisher/Invitrogen | #AM2295 |
| T4 polynucleotide kinase | NEB | #M0201S |
| FastAP thermosensitive alkaline phosphatase | Thermo Fisher Scientific | #EF0651 |
| T4 RNA ligase 1, high concentration | NEB | #M0437M |
| Proteinase K | NEB | #P8107S |
| SuperScript IV reverse transcriptase | Invitrogen | #18090050 |
| RNase inhibitor | NEB | #M0314L |
| RNase inhibitor | Produced in-house | N/A |

Kits

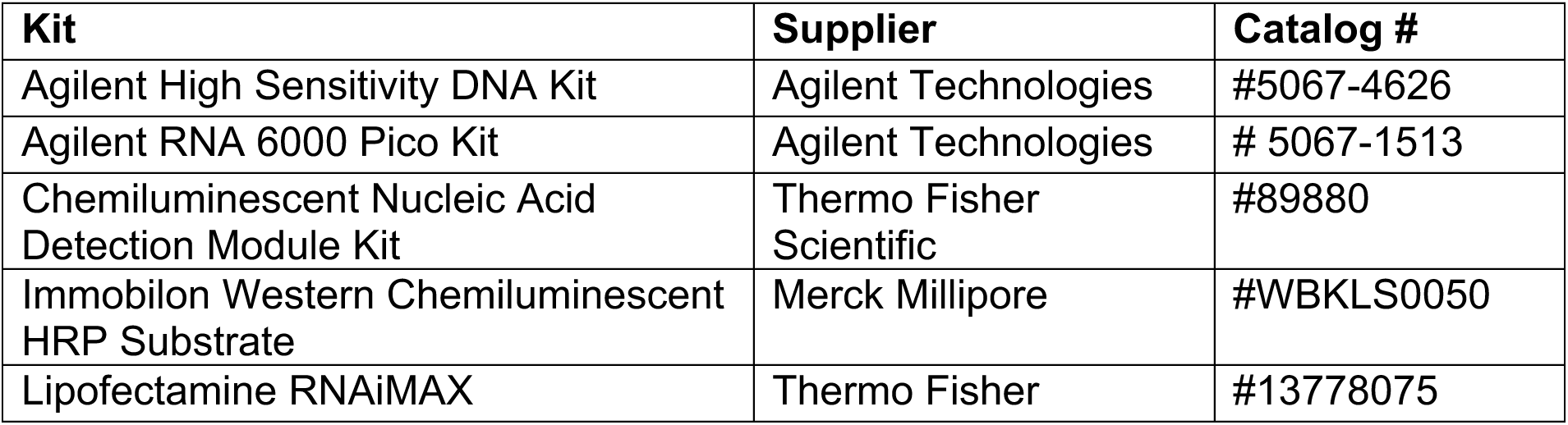
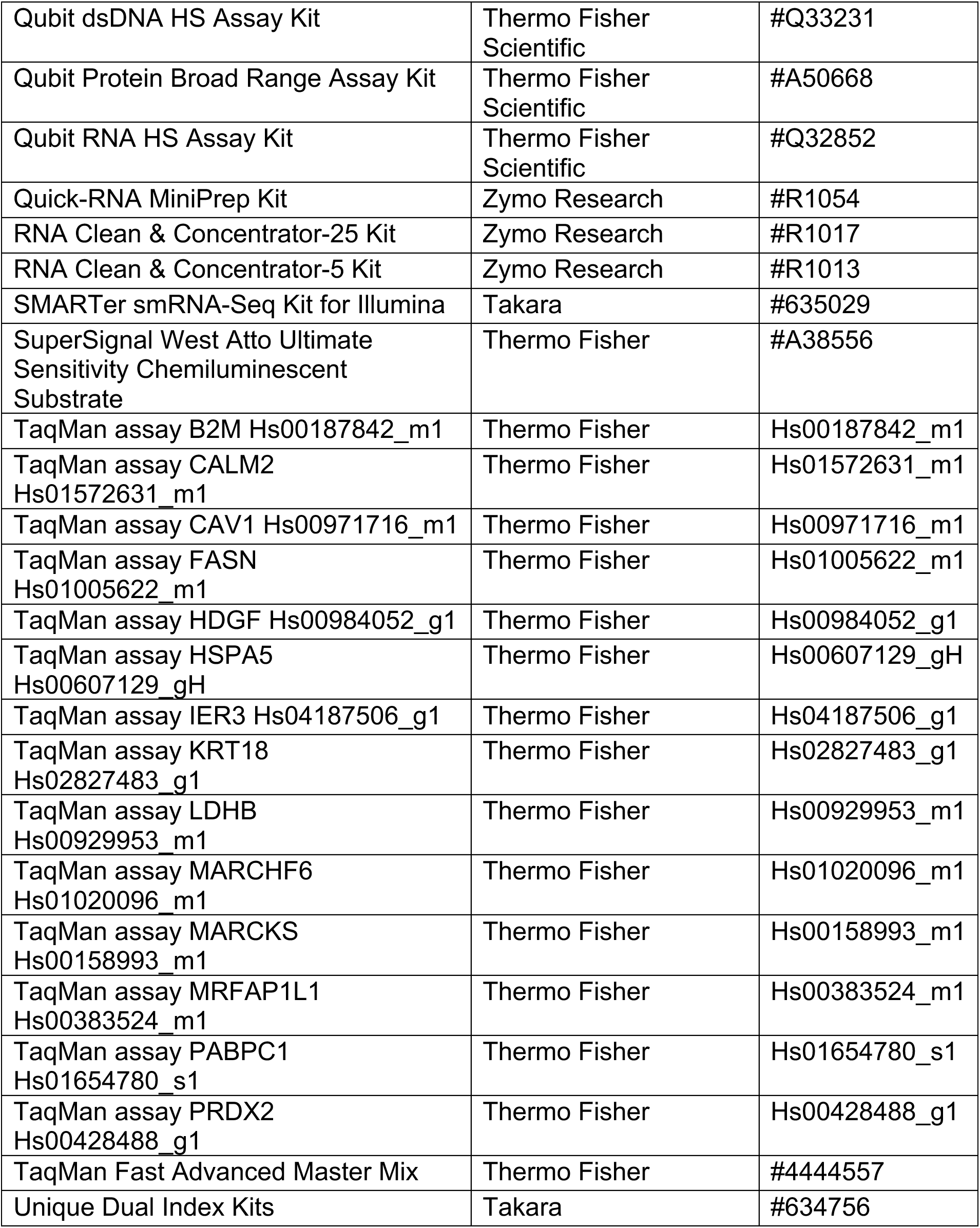

Reagents

| Reagent | Supplier | Catalog # |
| --- | --- | --- |
| Acid phenol/chloroform/isoamyl alcohol | Sigma | #P3803-400ML |
| Agencourt AMPure XP beads | Beckman Coulter | #A63880 |
| 4–15% Criterion™ TGX™ Precast Midi Protein Gel, 18 well, 30 µl | Bio-Rad | #5671084 |
| Dynabeads™ MyOne™ SILANE | Thermo Fisher Scientific | #37002D |
| Dynabeads™ Protein A | Thermo Fisher Scientific | #10002D |
| GlycoBlue | Invitrogen/Thermo Fisher Scientific | #AM9516 |
| NuPAGE 4× LDS sample loading buffer | Thermo Fisher | #NP0007 |
| pCp-biotin | Jena Bioscience | #NU-1706-BIO |
| Phase Lock Gel TM – Heavy | 5PRIME | #2302830 |
| Trans-Blot Turbo Midi Nitrocellulose Transfer Packs | Bio-Rad | #1704159 |
| TRIzol LS Reagent | Thermo Fisher Scientific | #10296028 |
| TURBO DNA-free Inactivation Reagent | Invitrogen | #AM1907 |

Proteins

Recombinant human PKM2 produced in-house (see Material and Methods)

Oligonucleotides and Plasmids

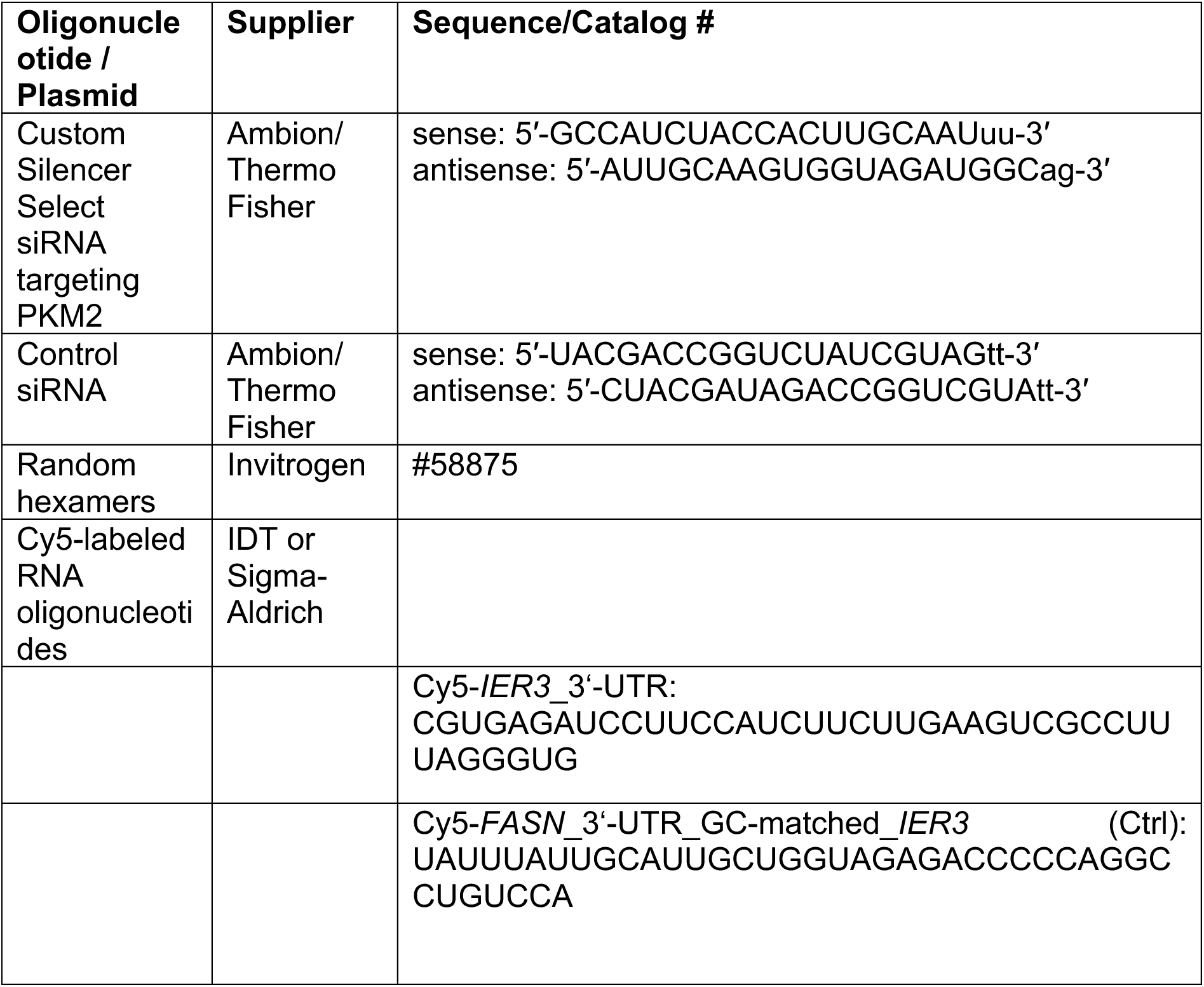
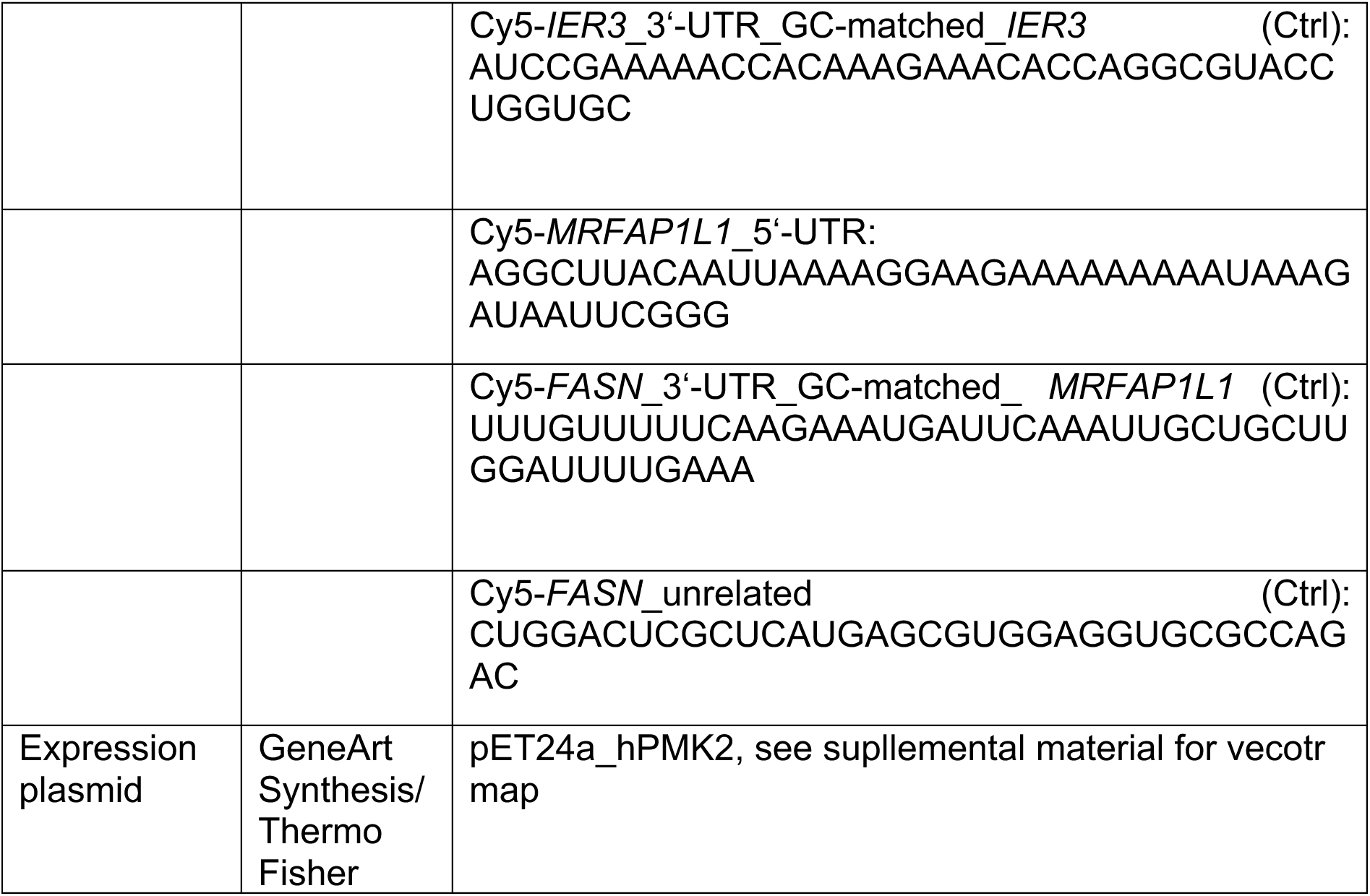

Data sources

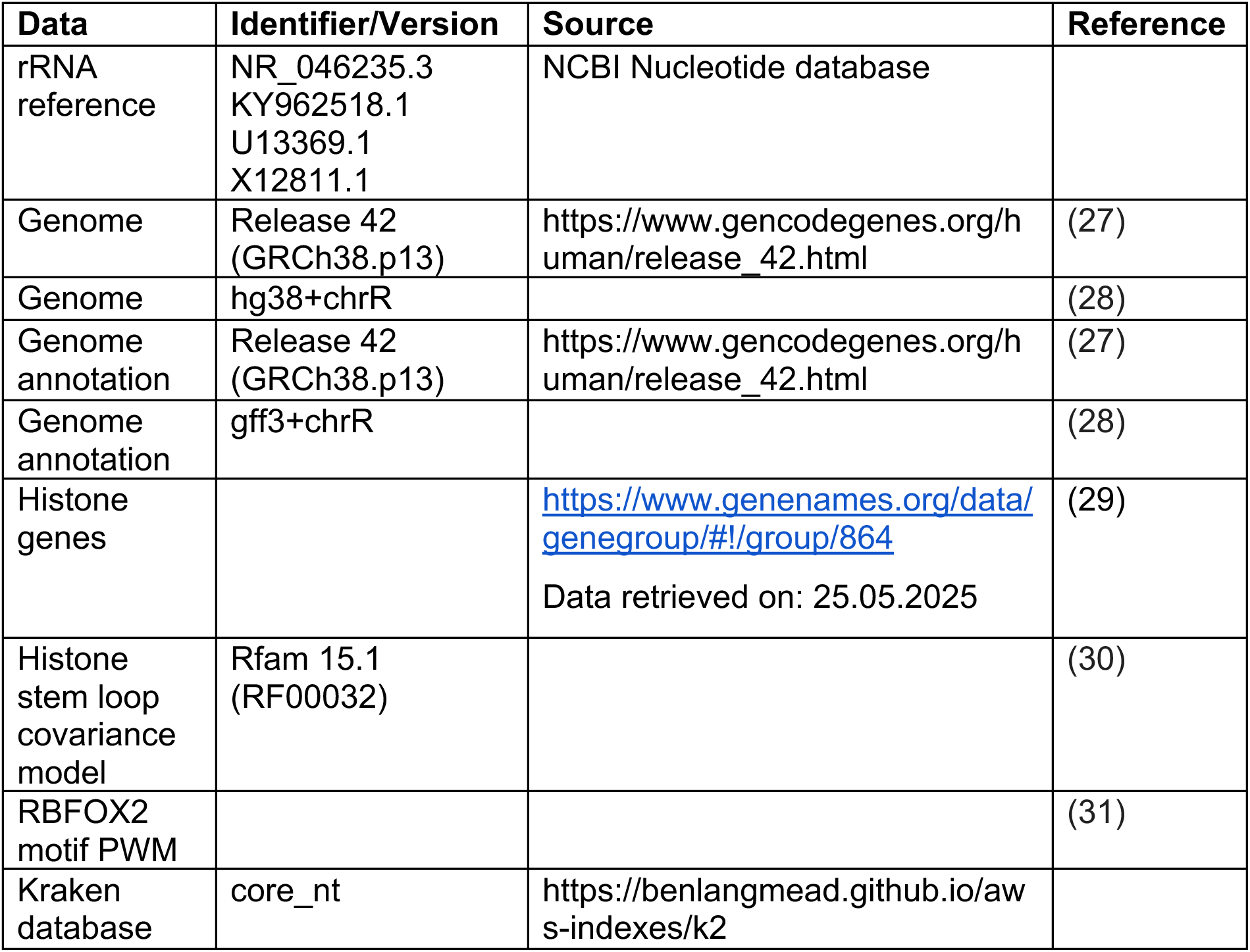

eCLIP data sources

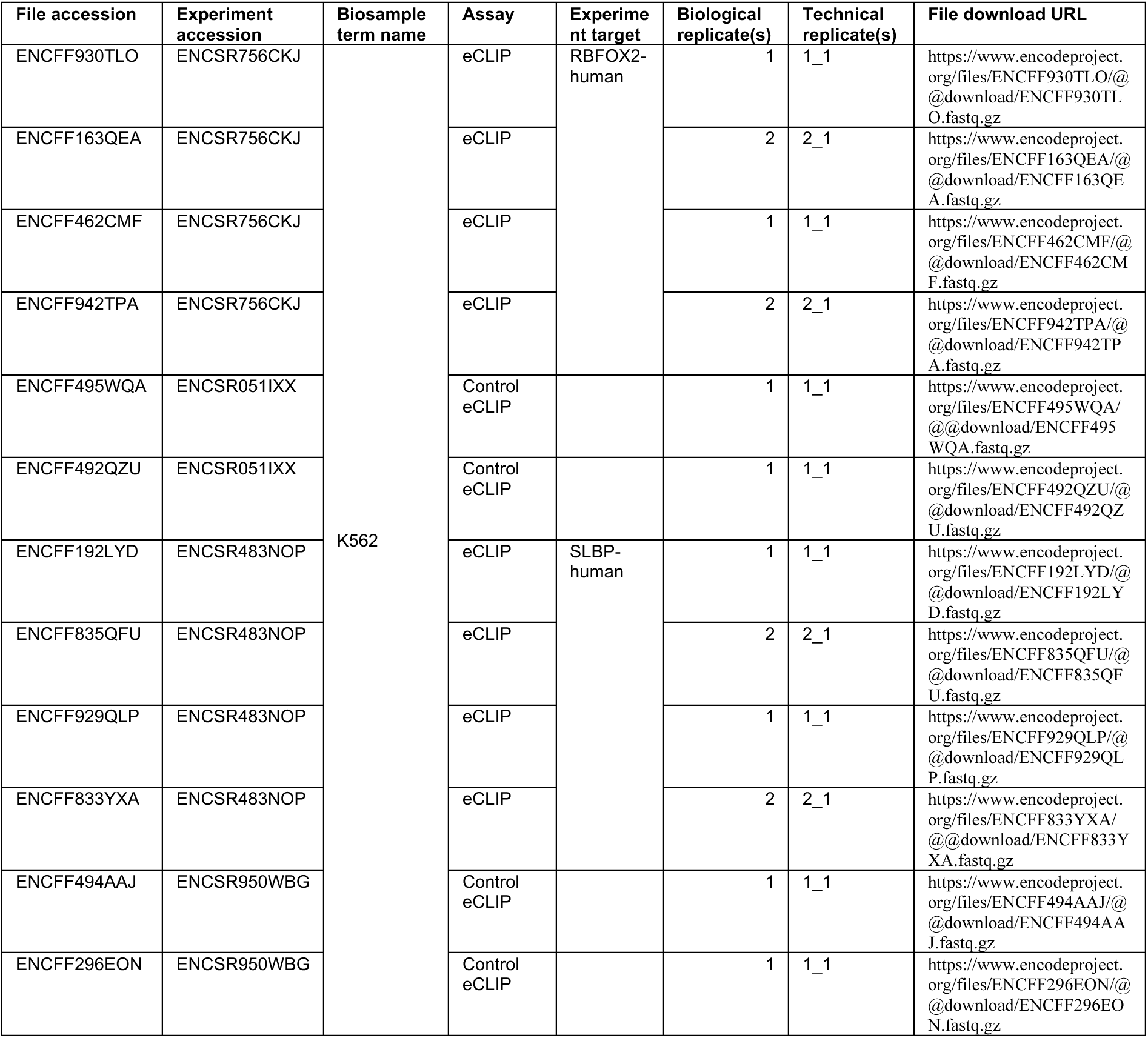

Bioinformatics tools

| Tool | Version(s) | Source | Reference |
| --- | --- | --- | --- |
| fastqc | 0.12.1 | https://www.bioinformatics.babraham.ac.uk/projects/fastqc/ |  |
| multiqc | 1.30 |  | (32) |
| seqkit | 2.13.0 |  | (33) |
| cutadapt | 5.1 |  | https://journal.embn et.org/index.php/em bnetjournal/article/view/200 |
| fastp | 1.0.1 |  | (34) |
| bbduk | 39.06, 39.08 |  | (35) |
| STAR | 2.7.10b |  | (36) |
| samtools | 1.21 |  | (37) |
| Kraken2<br>KrakenTools | 1.2.1 |  | (38,39) |
| Shoji | 0.44.3 | https://github.com/EMBL-Hentze-group/Shoji |  |
| UMI-tools | 1.1.6 |  | (40) |
| bedtools | 2.31.1 |  | (41) |
| UCSC-bedGrapToBigWig<br>UCSC-bedSort | 469 | https://hgdownload.soe.ucsc.edu/admin/exe/ |  |
| sourmash | 4.8.14 |  | https://joss.theoj.org/papers/10.21105/joss.06830 |
| ngs-statter | 1.20.3 | https://github.com/EMBL-Hentze-group/ngs-statter |  |
| JBrowse 2 | 2.10.1 - 3.6.5 |  | (42) |
| meme-suite | 5.5.4 |  | (43) |
| NextFlow | 25.04, 25.10 |  | (44) |
| Infernal/ cmscan | 1.1.5 |  | (45) |
| R | 4.5.1 | https://www.r-project.org/ |  |
| DEWSeq | 1.22.0 |  | (46) |
| DEWSeqHelper | 0.4.1 | https://github.com/EMBL-Hentze-group/dewseq_helper |  |
| DESeq2 | 1.48.2 |  | (47) |
| ZINBWAVE | 1.30 |  | (48) |
| GenomicRanges | 1.60 |  | (49) |
| IHW | 1.36 |  | (50) |
| tidyverse | 2.0 | https://tidyverse.org/ |  |
| pheatmap | 1.0.13 | https://raivokolde.r-universe.dev/pheatmap |  |
| ggrepel | 0.9.6 | https://ggrepel.slowkow.com/ |  |

BIOLOGICAL RESOURCES

| Reagent | Supplier | Catalog # |
| --- | --- | --- |
| HeLa | cells were maintained from long-standing laboratory stocks;<br>cell line authentication was performed | N/A |
| K562 | cells were maintained from long-standing laboratory stocks;<br>cell line authentication was performed | N/A |

## METHODS

### Cell Culture

HeLa cells were cultured in high-glucose (4.5 g/L D-glucose) Dulbecco’s modified Eagle’s medium (DMEM) supplemented with 2 mM L-glutamine, 10% heat-inactivated fetal bovine serum (FBS) (10270106, Thermo Fisher Scientific) and 100 U/mL penicillin-streptomycin in an incubator at 37 °C and 5% CO_2_.

K562 cells were cultured in Roswell Park Memorial Institute (RPMI) medium supplemented with 2 mM L-glutamine, 10% heat-inactivated fetal bovine serum (FBS) and 100 U/mL penicillin-streptomycin in an incubator at 37 °C and 5% CO_2_ in suspension cell culture dishes.

### Complex Capture (2C)

#### Generation of cross-linked HeLa cell lysates for 2C analysis

HeLa cells were cultured in 15 cm culture dishes until 80-90% confluent. Plates were placed on ice and washed twice with ice-cold PBS. Remaining PBS was removed completely and plates were left to dry upside down for 30 s. While on ice, cells were cross-linked using the SpectroLinker™ XL-1500 UV-Crosslinker (150 mJ/cm^2^ UV radiation). At least one plate per condition was kept as the non-cross-linked control. Immediately after cross-linking, cells were lysed and scraped in 500 µL of ice-cold iCLIP100 lysis buffer (see Buffers, supplemental material) per plate. Lysates were vortexed every 5 min for 15 min and kept on ice in between and subsequently sonicated using a Diagenode sonicator Bioruptor® Plus (5 cycles; 30 seconds ON, 30 seconds OFF; high power mode at 4 °C). Samples were centrifuged at 13,400 g for 10 min at 4 °C. Supernatants were transferred to fresh tubes. Protein concentration was measured using the Qubit™ Protein Broad Range (BR) Assay Kit on a Qubit™ 4 Fluorometer.

#### Generation of cross-linked K562 lysates for 2C analysis

K562 cells were cultured until 5e5/ml were reached. Cells from one 10 cm dish were washed with ice-cold PBS and resuspended in 3 ml ice-cold PBS. The 3 ml cell suspension was spread on a fresh 10 cm suspension cell culture plate and cross-linked on ice as described for HeLa cells. Cells were scraped and the suspension was centrifuged at 200 g for 3 min. The cell pellet was washed with 10 ml ice-cold PBS. The cell pellet was resuspended in 100 µl iCLIP100 lysis buffer (see Buffers, supplemental material) and further processed as described for HeLa cells.

#### 2C procedure

2C procedure was adapted from (51). 1.5 mg of protein lysate were diluted to a total volume of 300 µL in iCLIP100 lysis buffer (see Buffers, supplemental material). RNA extraction was performed using the Quick-RNA™ MiniPrep Kit from Zymo Research utilizing a vacuum manifold connected to a vacuum pump and following the manufacturer’s instructions. For elution, the columns were transferred to fresh collection tubes and residual RNA wash buffer was removed by centrifugation at 12,000 g for 1 min at RT. After transferring the columns to fresh microreaction tubes, 100 µL of nuclease-free water pre-warmed to 50 °C was added on top of each column matrix and incubated at 50 °C for 5 min. RNA was eluted by centrifugation at 12,000 g for 1 min at RT. The RNA yield was quantified using a NanoDrop 1000 spectrophotometer. 95 µL of each RNA extract were subjected to DNase-treatment using 2.5 µL of Turbo™ DNase (2 U/µL) (Thermo Fisher) and 10 µL of 10X Turbo™ DNase buffer. Of each set, one sample was treated with 5 µL of Ambion RNase I (Invitrogen) (100 U/µL) (negative control). Samples were incubated at 37 °C for 30 min, 1200 rpm shaking.

A second round of column purification was performed using the RNA Clean & Concentrator-25 kit by Zymo Research following manufacturer’s instructions and utilizing a vacuum manifold attached to a vacuum pump. For elution, the columns were transferred to fresh collection tubes and residual RNA wash buffer was removed by centrifugation at 12,000 g for 1 min at RT. After transferring the columns to fresh microreaction tubes, 25 µL of nuclease-free water pre-warmed to 50 °C was added on top of each column matrix and incubated at 50 °C for 5 min. The final RNA yield of the flowthrough was quantified using a NanoDrop 1000 spectrophotometer.

An equivalent amount of RNA was treated with 5 µL of Ambion RNase I (Invitrogen) (100 U/µL) or 5 µL of nuclease-free water, respectively, and incubated at 37 °C for 30 min, 1200 rpm shaking. 6.5 µL of freshly prepared 4X NuPAGE lithium dodecyl sulfate (LDS) sample loading buffer (Thermo Fisher) supplemented with 285 mM dithiothreitol (DTT) were added to each sample and input controls and incubated at 70 °C for 10 min, 1200 rpm shaking. 30 µL of prepared sample or input control were loaded onto 4-15% Criterion™ TGX™ Precast Midi Protein Gels (BioRad) alongside 5 µL of BioRad Precision Plus Protein Dual Color Standards (BioRad). The gels were run at 200 V for approximately 30 min. Protein transfer was performed using a TransBlot™ Turbo Transfer System (BioRad) on mixed molecular weight turbo blotting mode with Trans-Blot™ Turbo Midi 0.2 µm Nitrocellulose Transfer Packs (BioRad). Membranes were blocked in 5% milk in 1x tris-buffered saline with Tween^®^ 20 (TBST) for 30 min at RT followed by three quick TBST washes. Western Blotting was performed according to the BioRad protocol. The following primary and secondary antibodies were used for western blot analyses: ACTIN (AC-15 Sigma-Aldrich, 1:10000 in 5% Milk in TBST), PKM2 (D78A4 Cell Signaling, 1:1000 in 5% BSA in TBST), HuR (#11910-1-AP Proteintech, 1:2000 in 5% Milk in TBST), histone-H3 (#17168-1-AP Proteintech, 1:1000 in 5% Milk in TBST), HRP-conjugated anti-mouse IgG (#AB6789, Abcam, 1:5000 in 5% Milk in TBST), HRP-conjugated anti-rabbit IgG (AB97051, Abcam, 1:5000 in 5% Milk in TBST). Blots were developed with Immobilon^®^ Western Chemiluminescent HRP Substrate (Merck Millipore) or SuperSignal™ West Atto Ultimate Sensitivity Chemiluminescent Substrate (Thermo Fisher). Western blot signal detection was carried out on a ChemiDoc Go Imaging System (BioRad).

### T4 Polynucleotide Kinase (PNK) Assay

Lysates were generated as described under *Complex Capture (2C)* using PNK lysis buffer (see Buffers, supplemental material). Protein A magnetic beads (Dynabeads™, Thermo Fisher) were used for immunoprecipitation of rabbit antibodies. For each IP reaction, 50 µL of beads were used. Beads were washed three times with 1 mL of PNK lysis buffer, followed by resuspension in 100 µL lysis buffer per IP. The beads were distributed into new microreaction tubes and the volume was adjusted to 400 µL with PNK lysis buffer. Antibodies were added as follows: PKM2 (D78A4 Cell Signaling, 1 µg/IP), rabbit IgG (Invitrogen #10500C, 1 µg/IP). Beads and antibodies were incubated on a rotating wheel at 13 rpm 4 °C for 2 h or overnight.

Cell lysates (500 µg total protein/condition) were adjusted to 500 µL using PNK lysis buffer (see Buffers, supplemental material). RNase digestion was performed using Ambion RNase I (Invitrogen) (1 U/mL, 2 U/mL, 4 U/mL, 8 U/mL) together with 2 µL Turbo DNase (Invitrogen). Digestions were carried out at 37 °C for 15 min at 850 rpm. Subsequently, 11 µL RNase inhibitor (produced in house) was added per sample. A 30 µL aliquot was reserved as input control. Following bead-antibody coupling, beads were washed three times with 1 mL PNK lysis buffer and resuspended in 50 µL PNK lysis buffer per IP. Beads were added to each digested lysate and incubated at 4 °C for 3 h on a rotating wheel.

Beads were washed three times with 1 mL lysis buffer (rotation at 13 rpm for 3 min at RT), followed by two washes with PNK high-salt buffer (see Buffers, supplemental material) under the same conditions. A final wash was performed three times with 1 mL PNK wash buffer (see Buffers, supplemental material). After the last wash, supernatants were carefully removed to ensure complete clearance. All post-IP handling was conducted in a radioactive-safe lab area. For the T4 polynucleotide kinase (PNK) reaction, the following master mix was prepared per reaction: 24 µL RNase-free water, 3 µL 10× PNK buffer, 8 µL T4 PNK enzyme (NEB). To initiate 5′-end labeling, 0.3 µL of [γ-32P]ATP (1 µCi/µL) (Hartmann Analytic) was added to each reaction. Beads were magnetically separated and resuspended in 30 µL of hot PNK reaction mix, incubated at 37 °C for 15 min at 850 rpm. Beads were washed four times with 1 mL PNK wash buffer. Labeled complexes were eluted by resuspending beads in 25 µL of 0.1 M glycine (pH 2.0), incubating for 5 min at RT, and neutralizing with 3.75 µL 1.5 M Tris-HCl (pH 8.5). The supernatant was recovered after magnetic separation.

Eluted samples were mixed with 8 µL of 4X NuPAGE LDS sample loading buffer (Thermo Fisher) supplemented with 285 mM DTT and incubated at 70 °C for 15 min. Prepared sample and input controls were loaded onto 4-15% Criterion™ TGX™ Precast Midi Protein Gels (BioRad) alongside 5 µL of BioRad Precision Plus Protein Dual Color Standards (BioRad) and run at 180 V for 45 minutes. Protein transfer was performed using a TransBlot™ Turbo Transfer System (BioRad) on mixed molecular weight turbo blotting mode with Trans-Blot™ Turbo Midi 0.2 µm Nitrocellulose Transfer Packs (BioRad). Membranes were rinsed in PBS and transferred to autoradiographic film. Films were exposed overnight or up to 48 h before analysis using a Typhoon™ laser scanner. Following exposure, western blot was performed as described under *Complex Capture (2C)*.

### pCp-Assay

Cross-linked lysates were generated as described under *Complex Capture (2C)*. Protein A magnetic beads (Dynabeads™, Thermo Fisher) were used for immunoprecipitation of rabbit antibodies. For each IP reaction, 20 µL of beads were used. Beads were washed three times with 1 mL of iCLIP lysis buffer (see Buffers, supplemental material), followed by resuspension in 100 µL lysis buffer per IP. Antibodies added: PKM2 (D78A4 Cell Signaling, 1 µg/IP), rabbit IgG (Invitrogen #10500C, 1 µg/IP). Beads and antibodies were incubated on a rotating wheel at 13 rpm in a cold room (4 °C) for 2 h or overnight. Cell lysates (500 µg total protein per sample) were adjusted to 500 µL using iCLIP lysis buffer. RNase digestion was performed using Ambion RNase I (Invitrogen) together with 2 µL Turbo DNase (Thermo Fisher). Digestions were carried out at 37 °C for 5 min at 850 rpm. Subsequently, 11 µL RNase inhibitor (produced in house) was added per sample. Alternatively, DNase digestion was performed and RNA was fragmented by sonication using a Diagenode Bioruptor^®^ Plus (20 cycles, 30 s ON / OFF, high power mode, 4 °C). A 5 µL aliquot was reserved as input control. Following bead-antibody coupling, beads were washed three times with 1 mL iCLIP lysis buffer and resuspended in 50 µL iCLIP lysis buffer per IP. Beads were added to each digested lysate and incubated at 4 °C for 2 h on a rotating wheel. Each sample was washed twice with 900 µL of ice-cold iCLIP100 lysis buffer, twice with 900 µL of ice-cold high salt wash buffer (see Buffers, supplemental material) without and once with LiCl (see Buffers, supplemental material) and once with 500 µL of ice-cold wash buffer (see Buffers, supplemental material) (rotating, 3 min per wash step). Beads were transferred to a fresh tube in ice-cold wash buffer. While separated on the magnetic rack, the samples were resuspended in 500 µL of ice-cold wash buffer, to which 500 µL of FastAP buffer (see Buffers, supplemental material) were added. After removing the supernatant, each sample was washed once with 500 µL of FastAP buffer and incubated with 50 µL of freshly prepared alkaline phosphatase reaction mix consisting of 5 µL 10X FastAP buffer, 2 µL murine RNase inhibitor (NEB), 2 µL Turbo™ DNase (Invitrogen), 3 µL FastAP™ thermosensitive alkaline phosphatase (Thermo Fisher) (1 U/µL) and 38 µL nuclease-free water at 37 °C for 15 min, 1200 rpm shaking. Without removing the alkaline phosphatase reaction mix, each sample was topped up and incubated with 50 µL of freshly prepared T4 polynucleotide kinase (PNK) reaction mix consisting of 20 µL 5X PNK buffer (pH6.5) (see Buffers, supplemental material), 2 µL T4 polynucleotide kinase (10 U/µL) (NEB) and 28 µL nuclease-free water at 37 °C for 20 min (interval shaking for 30 s, every 2 min). Reaction mixes were removed and each sample was washed in quick succession once with 500 µL of ice-cold wash buffer, twice with 500 µL of ice-cold high salt wash buffer and twice with 500 µL of ice-cold wash buffer.

On-bead ligation of the immunoprecipitated, cross-linked RNA was performed by adding 19 µL of freshly prepared biotinylation reaction mix consisting of 0.5 µL 1% (v/v) Tween^®^20, 2 µL 10X T4 RNA ligation buffer without DTT (see Buffers, supplemental material), 0.7 µL 100% dimethyl sulfoxide (DMSO), 0.2 µL 100 mM ATP (NEB), 0.5 µL Cytidine-5’-phosphate-3’-(6-aminohexyl)phosphate labelled with biotin (pCp-biotin) (Jena Bioscience), 0.5 µL RNase inhibitor (NEB), 8 µL polyethylene glycol (PEG) 8000 and 6.6 µL nuclease-free water. 1 µL of high-concentration T4 RNA ligase 1 (30 U/µL) (NEB) was added per sample and incubated at 16 °C for 2 h.

Without removing the biotinylation reaction mix, the samples were washed in quick succession twice with 500 µL of ice-cold high salt wash buffer and three more times with 500 µL of ice-cold wash buffer. The wash buffer was completely removed and each sample was resuspended in 20 µL of wash buffer. Input controls were topped to a total volume of 20 µL with wash buffer. All samples and inputs were denatured and removed from the beads by the addition of 10.5 µL of 4X NuPAGE LDS sample loading buffer (Thermo Fisher) supplemented with 285 mM DTT, 70 °C for 10 min 1200 rpm shaking.

5 µL or 25 µL of prepared sample were loaded per well on two separate 4-15% Criterion™ TGX™ Precast Midi Protein Gels (BioRad) and run at 200 V for approximately 30 min. Transfer was performed using a TransBlot™ Turbo Transfer System (BioRad) on mixed molecular weight turbo blotting mode with Trans-Blot™ Turbo Midi 0.2 µm Nitrocellulose Transfer Packs (BioRad). From this point onwards, the two membranes were processed independently.

To detect protein signals, the low input membrane (5 µl sample) was used. Blocking, antibody incubation and development were performed as described under *Complex Capture* (2C). As secondary antibody HRP-conjugated anti-rabbit IgG, light chain specific (#SA00001-7L Proteintech, 1:5,000 in 3% Milk-TBST) was used.

To detect RNA cross-linked to protein, the high input membrane (25 µl sample) was used. RNA was detected using the Chemiluminescent Nucleic Acid Detection Module Kit (Thermo Fisher Scientific) following manufacturer’s instructions. Detection was carried out on a ChemiDoc Go Imaging System (BioRad).

### Generation of PKM2 knockdown (KD) samples

Reverse transfection in 6-well format was carried out using Lipofecatmine^TM^ RNAiMax (Thermo Fisher) following manufacturer’s instructions (150,000 cells/well, 30 pMol RNA/well, 4 µl Lipofectamine/well). A custom Silencer Select® siRNA targeting PKM2 (Sense: 5’-GCCAUCUACCACUUGCAAUUU, Antisense: 5’-AUUGCAAGUGGUAGAUGGCag-3’) and control siRNA were acquired from Ambion (now Thermo Fisher). Medium was exchanged after 24 h and cell lysates were generated after 72 h as described under *Complex Capture (2C)* using 100 µl of iCLIP lysis buffer/well. Subsequently, Western Blot analysis of KD efficiency and pCp Assay were performed (see *pCp-Assay*).

### Sonication for RNA Fragmentation

HeLa or K562 lysates were used (see *Complex Capture (2C)* for preparation). 0.2 µg/µl of protein in 50 µl iCLIP 100 buffer were used per condition and supplemented with 0.2 µl Turbo DNase (Thermo Fisher). Controls were treated with Ambion RNase I (Invitrogen) (80 U/ml). Samples were incubated at 37 °C for 5 min, 1200 rpm shaking. Sonication samples were sonicated in a Diagenode Bioruptor™ Plus sonicator (0 - 20 cycles; 30 seconds ON, 30 seconds OFF; high power mode at 4 °C). For comparability with CLIP approaches, proteins were digested using 127 μL ProK digestion solution per sample (11 μL ProK (NEB cat # P8107B), 100 mM Tris pH 7.5, 50 mM NaCl, 10 mM EDTA, 0.2% SDS) at 37°C for 5 min and 50°C for 20 min with interval mixing. ProK digestion solution was preincubated at 37°C for 20 min to inactivate RNases.

Total RNA was extracted from samples using TRIzol™ LS Reagent (Thermo Fisher Scientific) according to the manufacturer’s protocol with slight modifications. Briefly, 360 µL of TRIzol LS reagent was added to the samples and incubated for 5 min at RT. Subsequently, 96 µL of chloroform was added, and the mixture was vigorously shaken by hand for 15 s. After a 3-min incubation at RT, samples were centrifuged at 12,000 × g for 15 min at 4 °C to separate the phases. The aqueous phase was carefully transferred to a new tube, to which 240 µL of isopropanol and 0.6 µL of GlycoBlue™ (Invitrogen) were added to facilitate RNA precipitation. Samples were then incubated ON or alternatively for 30 min in a pre-chilled metal rack at −80 °C. RNA was pelleted by centrifugation at 13,000 rpm for 1 h at 4 °C. The supernatant was discarded, and the pellet was washed with 100 µL of freshly prepared 70% ethanol. After centrifugation at 13,000 rpm for 15 min at 4 °C, ethanol was removed. The pellet was air-dried for 1-2 min. RNA was resuspended in 8 µL of RNase-free water. RNA fragment length was assessed by running the RNA 6000 Pico Kit (Agilent Technologies) on a 2100 Bioanalyzer (Agilent Technologies) following manufacturer’s instructions.

### soniCLIP Protocol

#### Optimization of Immunoprecipitation

IP was performed as described under *pCp-Assay*. For optimization purposes different amounts of antibody were tested. Antibodies tested: PKM2 (D78A4 Cell Signaling; 0.6, 1, 2 µg), SLBP (ThermoFisher A303-968A; 1, 3, 6 µg), RBFOX2 (ThermoFisher A300-864A; 1, 3, 6 µg), rabbit IgG (Invitrogen #10500C). In the PKM2 IP assay the effects of an additional washing step with LiCl high salt wash buffer were investigated and subsequently included in all IPs. After IP, efficiency was assessed by western blot as described under *Complex Capture (2C)*.

Lysates of HeLa and K562 cells subjected to UV crosslinking were generated as described under *Complex Capture (2C)*, which includes a first round of 5 min sonication.

Protein A magnetic beads (Dynabeads™, Thermo Fisher) were used for immunoprecipitation of rabbit antibodies. For each IP reaction, 20 µL of beads were used. Beads were washed three times with 1 mL of iCLIP lysis buffer (see Buffers, supplemental material), followed by resuspension in 100 µL lysis buffer per IP. Antibodies added: PKM2 (D78A4 Cell Signaling, 2 µg/IP), SLBP (ThermoFisher A303-968A; 6 µg/IP) or RBFOX2 (ThermoFisher A300-864A; 3 µg/IP). Beads and antibodies were incubated on a rotating wheel at 13 rpm in a cold room (4 °C) for 2 h or overnight. For preclearing, 20 μL of Protein A magnetic beads (Dynabeads, Thermo Fisher Scientific) per IP was washed twice in iCLIP100 buffer (see Buffers, supplemental material), resuspended to 50 μL, and added to thawed HeLa cell lysates (500 μg protein in 500 μL iCLIP100 buffer). After 30 min rotation at 4 °C, beads were removed magnetically and supernatants transferred to fresh tubes.

DNase digestion was performed using 2 µL Turbo DNase (Thermo Fisher) at 37 °C for 5 min at 850 rpm. Subsequently, 1 µL RNase inhibitor (NEB) was added per sample. RNA was fragmented by sonication using a Diagenode Bioruptor^®^ Plus (20 cycles (HeLa) or 15 cycles (K562), 30 s ON / OFF, high power mode, 4 °C). A 10 µL aliquot (2%) was reserved as input control after sonication. Following bead-antibody coupling, beads were washed three times with 1 mL iCLIP lysis buffer and resuspended in 50 µL iCLIP lysis buffer per IP. Beads were added to each fragmented lysate and incubated at 4 °C for 2 h on a rotating wheel. Biological quadruplicates of HeLa or K562 cell lysates were processed, respectively. During the incubation time of the immunoprecipitation reaction, the input samples were processed.

Input RNA (10 µl) was first treated with 15.6 μL FastAP master mix (10 μL nuclease-free water, 2.6 μL 10× FastAP buffer, 0.5 μL RNase inhibitor (NEB), 2.5 μL FastAP enzyme (Thermo Fisher Scientific)) and incubated at 37 °C for 15 min, 1200 rpm shaking. 75 μL PNK master mix (45 μL nuclease-free water, 20 μL 5× PNK buffer (pH 6.5) (see Buffers, supplemental material), 1 μL 0.1 M DTT, 1 μL Turbo™ DNase (Invitrogen), 1 μL RNase inhibitor (NEB), 7 μL PNK enzyme (NEB)) were added directly to the FastAP-treated RNA. Samples were incubated at 37 °C for 20 min with interval shaking (1000 rpm) every 2 min. RNA was purified using Dynabeads™ MyOne™ SILANE beads (Thermo Fisher Scientific). 20 µl beads per sample were washed once with 900 μL RLT buffer (Qiagen) and resuspended in 300 μL RLT. The slurry was added to the enzyme-treated RNA together with 10 μL of 5 M NaCl and 615 μL 100% ethanol, mixed, and rotated for 15 min at RT. Beads were washed once with 1 mL 75% ethanol, transferred to a fresh tube and washed twice more with 75% ethanol (30 s each). Beads were briefly spun (1000 rpm, 30 s), magnetized and residual liquid was removed. Beads were air-dried for 5 min and RNA was eluted in 10 μL nuclease-free water by incubating 5 min at RT. Input RNA was kept on ice until further processing. Following immunoprecipitation, each sample was washed twice with 900 µL of ice-cold iCLIP100 lysis buffer, twice with 900 µL of ice-cold high salt wash buffer (see Buffers, supplemental material) without and once with LiCl (see Buffers, supplemental material) and once with 500 µL of ice-cold wash buffer (see Buffers, supplemental material) (rotating, 3 min per wash step). Beads were transferred to a fresh tube in ice-cold wash buffer. While separated on the magnetic rack, the samples were resuspended in 500 µL of ice-cold wash buffer, to which 500 µL of FastAP buffer (see Buffers, supplemental material) were added. After removing the supernatant, each sample was washed once with 500 µL of FastAP buffer. FastAP buffer was removed and beads were incubated with 100 µL of freshly prepared alkaline phosphatase reaction mix consisting of 10 µL 10X FastAP buffer, 2 µL RNase inhibitor (NEB), 1 µL of Turbo™ DNase (Invitrogen), 8 µL of FastAP™ thermosensitive alkaline phosphatase (1 U/µL) (Thermo Fisher) and 79 µL of nuclease-free water at 37 °C for 15 min, 1200 rpm shaking. Without removing the alkaline phosphatase reaction mix, each sample was topped up and incubated with 300 µL of freshly prepared T4 polynucleotide kinase (PNK) reaction mix consisting of 60 µL 5X PNK buffer (pH6.5) (see Buffers, supplemental material), 7 µL T4 polynucleotide kinase (10 U/µL) (NEB), 3 µl 0.1M DTT, 1 µl RNAse Inhibitor (NEB), 8 µl Turbo™ DNase (Thermo Fisher) and 224 µL nuclease-free water at 37 °C for 20 min (interval shaking for 30 s every 2 min). Both reaction mixes were removed and each sample was washed in quick succession once with 500 µL of ice-cold wash buffer (see Buffers, supplemental material), twice with 500 µL of ice-cold high salt wash buffer (see Buffers, supplemental material) and twice with 500 µL of ice-cold wash buffer.

RNA was eluted from the beads by incubation with 127 μL Proteinase K (ProK) digestion solution (11 μL ProK (NEB, cat. #P8107B) in ProteinaseK buffer soniCLIP (see Buffers, supplemental material)). The digestion buffer was prewarmed for 20 min to 37 °C to inactivate residual RNases before being added to the bead-RNA complexes. Samples, including inputs, were incubated with 127 µl ProK digestion solution at 37 °C for 5 min, followed by 50 °C for 20 min with interval mixing. Beads were then placed on a magnetic stand and 120 μL of the supernatant was transferred to a clean tube for downstream processing. Input samples were not further purified.

RNA was extracted using TRIzol™ LS Reagent (Thermo Fisher Scientific) following manufacturer’s guidelines with minor modifications. Briefly, 360 μL TRIzol LS was added to each sample and incubated for 5 min at RT, followed by the addition of 96 μL chloroform. Samples were shaken vigorously by hand for 15 s, incubated for 3 min at RT, and centrifuged at 12,000 × g for 15 min at 4 °C. The aqueous phase was transferred to a new tube, mixed with 240 μL isopropanol and 0.6 μL GlycoBlue (Thermo Fisher Scientific), and precipitated either overnight at −80 °C or for 30 min at −80 °C in a pre-chilled metal rack. RNA was pelleted by centrifugation at 13,000 rpm for 1 h at 4 °C, the supernatant removed, and the pellet washed with 100 μL freshly prepared 70% ethanol. After centrifugation at 13,000 rpm for 15 min at 4 °C, residual ethanol was removed, the pellet was briefly air-dried (1-2 min) and resuspended in 10 μL nuclease-free water. A 2.5 μL aliquot was taken for quality control and the remaining RNA was stored at −80 °C. IP and Input RNA was assessed by running the RNA 6000 Pico Kit (Agilent Technologies) on a 2100 Bioanalyzer (Agilent Technologies) following manufacturer’s instructions. Input was quantified using the Qubit™ RNA HS Assay Kit on a Qubit™ 4 Fluorometer.

For library construction, the SMARTer smRNA-Seq Kit for Illumina (Takara) was used following manufacturer’s instructions with minor modifications. 7 µl of RNA were used for the IPs as starting material, for inputs 0.3 - 1 ng of RNA were used. In the PCR amplification step only 50% of the cDNA was used and filled up to 100% with H2O, the remaining half was stored at −80°C, serving as a backup. All libraries were amplified with 22 cycles. Indexing was done using the Unique Dual Index Kits (Takara) instead of the provided barcodes. Size selection was done using Agencourt AMPure XP beads (Beckman Coulter) following the protocol provided in the SMARTer smRNA-Seq Kit for Illumina manual. Library validation was done on a Bioanalyzer (Agilent Technologies) using the Agilent High Sensitivity DNA Kit (Agilent Technologies). A Qubit™ 4 Fluorometer with the Qubit™ dsDNA HS Assay Kit was used for quantification.

Sequencing was performed on a NextSeq2000 device (Illumina) following sequencing kit manufacturer’s instructions in single-end mode reading 100 nucleotides. 2.2 pM of library were loaded on the instrument.

### Bioinformatic data analysis

This section describes the bioinformatics resources and analysis steps used in this study. The details of the data sources and tools used are given in the Data Sources and Bioinformatics tools tables respectively and the first two subsections and the latter subsections below describe the data analysis steps.

### Computational steps

#### Extracting histone 3’UTR stem loop reference locations

Infernal cmscan tool was used to scan the human reference genome (GRCh38.p13) using the histone stem loop covariance model (RF00032) retrieved from Rfam database (version: 15.1). The first scan was performed using cmscan options ‘--noali --incT 25 --notrunc’ and the scan results were filtered to remove the stem loop locations identified on chromosomal scaffolds. This resulted in 159 histone stem loop chromosomal locations, which were further used as the primary histone 3’ UTR stem loop references. In a second relaxed scan, we used cmscan options ‘--noali --notrunc’, (but without ‘--incT 25’ parameter) to identify stem loop locations that would have been omitted from the first scan. The genomic locations in this result overlapping with the primary reference, (described in the step above) were filtered out and we used the resulting 193 histone stem loop genomic locations as extended stem loop references.

#### Extracting RBFOX2 motif locations

To extract RBFOX2 motif locations from the human reference genome (GRCh38.p13), we first extracted the sequences within annotated genes (Gencode release 42 genome annotation) in the genome using bedtools fastaFromBed tool. A background model was generated from these extracted sequences using MEME-suite fasta-get-markov tool using the option ‘-m 5’. In the next step, the MEME-suite fimo tool was used to scan the sequences generated in the previous step, using the RBFOX2 motif Position Weight Matrix in meme format (see Data sources subsection) and the background model described before. The fimo tool parameters used in this analysis were ‘--no-pgc --norc --thresh 0.001 --text --no-qvaluè and the overlapping motif locations from the resulting analysis were merged to generate 595,338 motif locations which were used in further downstream analysis.

#### eCLIP data retrieval and preprocessing

RBFOX2 and SLBP eCLIP datasets (generated by (4)) were retrieved from the ENCODE project portal. The details of the retrieved files are given in the Materials section. The fastq files provided by the ENCODE project portal were already demultiplexed, and the UMI sequences were appended to the beginning of the read identifiers. In this dataset, the read identifier lines were pre-processed to move UMI sequences to the end of the read identifier, for UMI deduplication using UMI-tools.

### Data analysis workflow

A custom Nextflow-based pipeline has been developed for analyzing soniCLIP and eCLIP data. The steps in this workflow are described below:

#### Quality control and adapter trimming

The quality of the sequences was assessed using a combination of FastQC and MultiQC tools at several steps (raw data, after adapter trimming, after rRNA removal and after contamination check) in this workflow. Additionally, we generated k-mer abundance signatures for each of the samples at every step using the sourmash suite. Computing cosine similarity between these signatures and clustering the similarity profiles using hierarchical clustering (as implemented in sourmash suite) served as an additional quality control step, enabling the visualization of sample-specific similarities, both within and between treatment groups.

For soniCLIP datasets (soniCLIP-benchmark and PKM2) polyA adapter at the 3’ end, Illumina universal adapters, polyX tails and low complexity reads were trimmed using the fastp tool. For eCLIP datasets we adopted the cutadapt based two-step trimming strategy described in (4).

The read length distribution per sample was estimated from the adapter-trimmed reads using the fx2tab function from seqkit, and the counts per read length per sample were aggregated and visualized as a line plot using ggplot2 for the soniCLIP-benchmark and PKM2 analysis.

#### rRNA-filtering and genome alignment

For the benchmark analyses (soniCLIP-benchmark and eCLIP-benchmark), trimmed reads were first aligned to rRNA sequences using bbduk, and rRNA-free reads were subsequently mapped to the human reference genome (GRCh38) using STAR. For the PKM2 analysis, we skipped the rRNA trimming step for mapping to a custom GRCh38 genome with an artificial ribosomal DNA chromosome (28) as the reference genome. For a more accurate mapping to novel splice junctions, we followed a 2-pass mapping strategy with STAR aligner, as described in (52). For eCLIP data, aligned reads were deduplicated using UMI-tools dedup function.

#### Contamination estimation

We incorporated a Kraken2 based contamination estimation to our data analysis pipeline to assess whether the datasets were contaminated with RNA from external sources, as reported by (12). In this step, the reads that did not map to the reference genome were extracted and classified as either known (contamination) or unknown based on the precomputed Kraken2 core_nt database. A combination of KrakenTools scripts (kreport2mpa.py, combine_mpa.py) and collect-reports function in ngs-statter package were used for the aggregation and visualization of read classification reports.

#### Post processing

Shoji annotation and createSlidingWindows functions were used to flatten gene annotation (GFF3) and to generate sliding (overlapping) genomic windows from the flattened annotations. Crosslink sites were extracted from genome-mapped reads using the shoji extract function and were summarized per genomic window using the shoji count function. Shoji createMatrix function was used to aggregate per-sample-level counts into three R-friendly tables: a window-level annotation table, a crosslink count table that summarized crosslink counts across all samples per genomic windows and a max count table which contained the highest single-nucleotide crosslink count per window per sample. In the statistical analysis step using DEWSeq, windows with low crosslink counts in the crosslink count table were filtered out using the max count table. The filtered crosslink count table was used for further statistical analysis.

#### Track generation

For all the datasets, crosslink sites (from Shoji extract step in post processing) were processed using bedtools-genomecov function and coordinate sorted with UCSC-bedSort to generate bedgraph files. These bedgraph files were then converted into BigWig files using UCSC-bedGraphToBigWig and visualized using a custom deployment of the JBrowse 2 server.

#### Summary statistics generation for bioinformatic data analysis

A combination of seqkit stats command and ngs-statter STAR command were used to generate read count / read alignment summary statistics from the various steps in the workflow. The ngs-statter sample-stats command was used to aggregate the read data statistics at different data processing steps per sample and all per-sample read data statistics were then merged into a final summary table using the ngs-statter compile-stats command. The merged table was processed and visualized using the ggplot2 package.

### Statistical analysis and normalization

For the soniCLIP-benchmark and PKM2 data, to account for zero inflation (excess zeros), we used ZINBWaVE package to compute zero-inflated count observational weights on the window-level crosslink count matrix. In all studies, library size normalization was performed using the DESeq2 poscounts size-factor estimator, as it is robust to many zero or low counts. The resulting window-level normalized counts, together with ZINBWaVE observational weights, were tested for differential enrichment using the Wald test (from DEWSeq package), and the Independent Hypothesis Weighting (IHW) procedure was used for multiple testing correction. For eCLIP benchmark data, the differential enrichment test was performed without using ZINBWaVE observational weights. Subsequently, study-specific Log_2_ fold change and p-adjusted value thresholds (described in the subsequent section) were used to filter differentially enriched windows and overlapping enriched windows were merged into regions using the extractRegions function from DEWSeq.

The normalized counts were transformed using the variance stabilizing transformation (vst) and visualized with the plotPCA function in the DESeq2 package for soniCLIP-benchmark and PKM2 analysis. The scatterplots displaying the Log_2_ fold change and -Log_10_ transformed p-adjusted values from differentially expressed windows/regions for the soniCLIP-benchmark, eCLIP-benchmark, and PKM2 analysis were generated using ggplot2.

### Study specific adaptations

This section includes the study-specific design for computing differentially expressed windows and the parameters used to filter significant windows for the analysis.

#### soniCLIP-benchmark

For the soniCLIP-benchmark, windows with less than 5 counts across at least 4 samples were filtered out using the filterCounts function from the DEWSeq package. From the filtered windows, the differentially enriched genomic windows were identified across the following contrasts: SLBP-IP versus Input, RBFOX2-IP versus Input, SLBP-IP versus RBFOX2-IP, and RBFOX2-IP versus SLBP-IP. For SLBP, windows with p-adjusted value ≤ 0.01 and Log_2_ fold change ≥ 2 that were significant and overlapping in both the SLBP-IP versus Input and SLBP-IP versus RBFOX2-IP comparisons were merged into contiguous regions. For RBFOX2, significant windows passing the same filtering thresholds described for SLBP, and overlapping in RBFOX2-IP versus Input (as defined in the benchmark design) were merged into contiguous regions.

#### PKM2 analysis

For the PKM2 analysis, windows with less than 5 counts across at least 4 samples were filtered out using the filterCounts function from the DEWSeq package. From the filtered windows, differentially enriched windows were identified for the following contrasts: SLBP-IP versus Input, PKM2-IP versus Input, SLBP-IP versus PKM2-IP, and PKM2-IP versus SLBP-IP. For PKM2, windows with p-adjusted value ≤ 0.1 and Log_2_ fold change ≥ 1 that were significant and overlapping between PKM2-IP versus Input and PKM2-IP versus SLBP-IP comparisons were merged into contiguous regions.

#### eCLIP-benchmark

For the eCLIP-benchmark, windows with less than 3 counts across at least 2 samples were filtered out using the filterCounts function from the DEWSeq package. From the filtered windows, the differentially enriched windows were identified in SLBP-IP samples relative to the size-matched input (SMI), and in RBFOX2-IP samples relative to SMI. For each comparison, overlapping significantly differentially enriched windows with p-adjusted value ≤ 0.1 and Log_2_ fold change ≥ 1 were merged into contiguous regions.

### Benchmark analysis

#### Histone 3’ UTR stem loop structure and RBFOX2 motif analysis

For enriched SLBP regions across all studies, we computed the fraction of enriched regions overlapping reference histone 3′ UTR stem loop locations (described in the section *Extracting histone 3′ UTR stem loop reference locations*) using the structureOverlap function from the DEWSeqHelper package. For enriched RBFOX2 regions, we used the motifDistance function from DEWSeqHelper package to identify regions harboring a known RBFOX2 motif (described in the section *Extracting RBFOX2 motif locations*) or, for regions lacking a known motif, the same tool was used to calculate the distance to the nearest known RBFOX2 motif.

#### Crosslink profile plot

Crosslink profile plots (Figure 3D) were generated using the crosslink sites described in the *Post processing* section and the histone stem loop reference locations described in the *Extracting histone 3′ UTR stem loop reference locations* section, respectively using ngs-statter crosslink-line-plot command, using the following options “--most-5prime –show-group-mean” CLIP-seq data submission accession numbers, and repositories for the data analysis pipeline, helper packages, and annotation data used are provided in the Data Availability section.

### RIP-qPCR Assay

#### Formaldehyde-based crosslinking

Cells were grown until 80-90% confluent. 2×15 cm dishes were processed per replicate. For crosslinking, 16% formaldehyde (FA) (15710, Electron Microscopy) stock was freshly diluted in 1× PBS to a final concentration of 0.1%. Culture medium was removed immediately before crosslinking, and 18 mL of 0.1% FA solution was added per plate. Plates were gently rocked at RT for 9 min. The crosslinking solution was aspirated and cells were washed twice with PBS at RT. Crosslinking was quenched by adding 20 mL of 0.125 M glycine and gently rocking for 5 min at RT. The quenching solution was removed and cells were washed three times with ice-cold PBS (1-2 min per wash). From this step onwards, all procedures were performed on ice. Cells were lysed directly on the plates in chilled RIPA buffer (see Buffers, supplemental material) (500 μL per 15 cm dish). Cells were scraped on ice, and lysates were transferred to microcentrifuge tubes. Lysates were incubated on ice for at least 10 min prior to sonication. Chromatin was sheared in a Diagenode Bioruptor™ Plus sonicator (10 cycles; 30 seconds ON, 30 seconds OFF; high power mode at 4 °C). Lysates were cleared by centrifugation at 10,000 × g for 15 min at 4°C. Supernatants were collected, and protein concentrations were determined using the Qubit™ Protein Broad Range (BR) Assay Kit on a Qubit™ 4 Fluorometer. Lysates were stored at −70 °C until further use.

#### RNA-Immunoprecipitation, RNA isolation and cDNA synthesis

20 µl Protein A magnetic beads (Dynabeads, Thermo Fisher) were prepared per IP, washed twice in 1 mL RIPA qPCR lysis buffer (see Buffers, supplemental material), and resuspended in 100 μL RIPA qPCR lysis buffer. Beads were incubated overnight at 4 °C with the respective antibodies (PKM2: D78A4 Cell Signaling, 2 µg/IP; rabbit IgG: Invitrogen #10500C, 2 µg/IP). For preclearing, 15 μL bead slurry per IP was washed twice in RIPA, resuspended to 50 μL and added to thawed HeLa cell lysates (300 μg protein in 500 μL RIPA). After 30 min rotation at 4 °C, beads were removed magnetically and supernatants transferred to fresh tubes; 2% was reserved as input. Antibody-bead conjugates were washed thrice in RIPA, resuspended to 50 μL, and incubated with precleared lysates for 2 h at 4 °C, rotating. Beads were washed sequentially (3 min rotation, RT) with 1x RIPA qPCR lysis buffer, 2x RIPA high-salt (see Buffers, supplemental material), 1x RIPA LiCl high-salt (see Buffers, supplemental material), 2x RIP-qPCR wash buffer (see Buffers, supplemental material), and 1x RIPA, then transferred to fresh tubes.

Proteinase K treatment was performed in prewarmed PK mix (160 µl Proteinase K qPCR buffer (see Buffers, supplemental material); 20 µl Proteinase K (NEB)) (pre-incubated 15 min at 37 °C to inactivate RNases, supplemented with 1 µl RNase inhibitor (produced in house)). Inputs received 20 μL PK mix, IPs received 180 μL. Reactions were incubated at 55 °C for 30 min, 1200 rpm shaking. Supernatants were collected and input volumes adjusted to 180 μL.

Equal volumes of acid phenol/chloroform/isoamyl alcohol (pH 6.5) (P3803, Sigma) were added to each sample, mixed, and incubated at 37°C for 5 min at 1200 rpm shaking. Phase separation was performed using pre-spun Phase Lock Gel Heavy tubes (Prime) (12,000 rpm, 30 s, RT) followed by centrifugation at 13,000 × g for 15 min, RT. The aqueous phase was purified using the Zymo RNA Clean & Concentrator-5 kit according to the manufacturer’s total RNA >17 nt protocol. Briefly, two volumes RNA binding buffer and one volume ethanol were added before loading onto Zymo spin columns. After sequential washes with RNA Prep Buffer and RNA Wash Buffer, RNA was eluted twice with 10 μL pre-warmed (55°C) nuclease-free water, yielding ∼18 μL total and stored at −80°C.

DNase treatment was performed using Turbo™ DNase (Thermo Fisher Scientific) in 23.5 μL reactions containing 1 μL RNase inhibitor (NEB), 2.5 μL 10x buffer, and 2 μL Turbo™ DNase (Invitrogen) for 15 min at 37 °C. Reactions were stopped with 2.4 μL TURBO DNA-free™ Inactivation Reagent (Invitrogen), incubated 5 min at RT, centrifuged (10,000 g, 2 min) and supernatants (∼20 μL) were collected.

Reverse transcription was performed using SuperScript™ IV (Invitrogen) in 40 μL total volume. RNA (10 μL) was mixed with 2 μL random hexamers (#58875, Invitrogen, 150 ng), 2 μL dNTP mix, and nuclease-free water to 26 μL, heated to 65 °C for 5 min and chilled on ice. 14 µl RT master mix (8 μL 5x buffer, 2 μL 100 mM DTT, 2 μL water, 2 μL SSIV RT) was added; reactions were incubated at 23 °C for 10 min, 53 °C for 10 min and 80 °C for 10 min. The obtained cDNA was diluted 1:4 in RNase-free water and stored at −20 °C until further use.

#### qRT-PCR analysis

TaqMan Probes (Thermo Scientific) and TaqMan Fast Advanced Master Mix (ThermoFisher) were used on a QuantStudio 6 Flex Real-Time PCR System (Applied Biosystems). Analysis was performed as previously described (53,54), with IgG control samples serving as the background reference. For each qPCR assay, the Ct value from the RIP RNA fraction was first normalized to the corresponding Input RNA fraction Ct value (ΔCt) to account for differences in RNA sample preparation. The ΔCt was calculated as:

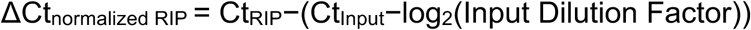

The percentage of Input RNA for each RIP fraction was calculated using the equation:

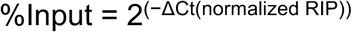

For background correction, the normalized RIP ΔCt values were adjusted by subtracting the mean ΔCt of the normalized IgG controls for the corresponding assay, yielding the first ΔΔCt:

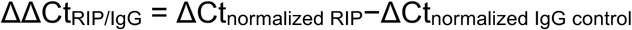

Finally, fold enrichment above the sample-specific background was determined by linear conversion of the ΔΔCt values:

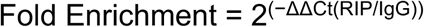

### PKM2 Protein Purification

Production and purification of recombinant PKM2 was performed by the EMBL Protein Production and Purification Core Facility. *Escherichia coli* BL21(DE3)-CodonPlus-RIL competent cells from Stratagene were freshly transformed with the expression plasmid (pET24a_hPMK2, Thermo fisher / GeneArt Synthesis, see Vector maps, supplemental material). Overnight (ON), precultures for large scale expression were grown at 37 °C in Lennox Broth medium supplemented with 30 μg/mL kanamycin and 34 μg/mL chloramphenicol. To inoculate the large-scale expression cultures, 10 mL of preculture were added to 1 L of Terrific Broth medium supplemented with 2 mM MgSO_4_, 0.05% glucose, 1.5% lactose, 30 μg/mL kanamycin and 34 μg/mL chloramphenicol. Cultures were grown at 37°C until optical density (OD) at 600 nm reached around 0.6, after which the growth temperature was reduced to 18 °C. Following ON auto-expression at 18 °C, the cultures were harvested by centrifugation at 5,000 g for 30 min at 4 °C and the pellets were flash-frozen in liquid nitrogen and stored at −80 °C until starting the protein purification.

The cell pellet was resuspended in ice-cold protein production lysis buffer (see Buffers, supplemental material). All further purification steps were performed at 4-8 °C. Cells were lysed by five passages through a microfluidizer, followed by centrifugation at 14,000 g for 20 min at 4 °C. The cleared lysate was loaded onto a 5 mL Protino nickel-nitrilotriacetic (Ni-NTA) acid fast protein liquid chromatography column by Macherey-Nagel pre-equilibrated with equilibration buffer (see Buffers, supplemental material). After loading, the Ni-NTA column was washed with equilibration buffer and eluted with equilibration buffer supplemented with 250 mM imidazole. After sodium dodecyl sulfate-polyacrylamide gel electrophoresis (SDS-PAGE) analysis, elution fractions containing PKM2 were pooled, the histidine-tag was cleaved by addition of histidine-tag specific tobacco etch virus (His-TEV) protease in a ratio of 1:100 of protein to protease and dialysed overnight at 4°C against dialysis buffer (see Buffers, supplemental material). The next day, cleaved PKM2 was further purified using a combination of anion exchange and heparin chromatography. The elution fractions from the Ni-NTA purification were diluted 6-fold with 50 mM HEPES (pH 7.4) supplemented with 10% glycerol to lower the NaCl concentration to 50 mM and then loaded onto a 5 mL HiTrap™ Q HP anion exchange chromatography column by Cytiva coupled in tandem to a 5 mL HiTrap™ Heparin HP affinity column by Cytiva. After washing with ion exchange equilibration buffer (see Buffers, supplemental material), the HiTrap Q HP anion exchange chromatography column was removed and PKM2 protein was eluted from the HiTrap Heparin HP column using a gradient ranging from 50 mM NaCl to 1 M NaCl over 20 column volumes. The elution fractions containing PKM2 were pooled, concentrated to 5 mL and injected into a HiLoad™ 16/600 Superdex™ 200 pg size-exclusion chromatography (SEC) column pre-equilibrated with SEC equilibration buffer (see Buffers, supplemental material). The retrieved PKM2 elution fractions were pooled and concentrated to 6-9 mg/mL. The final PKM2 protein was aliquoted, flash-frozen in liquid nitrogen and stored at −80°C until use.

The identity of the PKM2 protein was verified by mass spectrometry and its tetrameric oligomerization state and the absence of aggregates was confirmed by size exclusion chromatography with multi-angle light scattering. Appropriate stability in the storage buffer was assessed by NanoDSF analysis. PKM2 proteins were expressed and purified as independent biological triplicates.

### Electrophoretic mobility shift assay (EMSA)

A 20 nM solution of the respective Cy5-labelled RNA (see RNA oligos, supplemental material) in 2xD buffer (see Buffers, supplemental material) was denatured at 95 °C for 2.5 min, stabilized by the addition of MgCl_2_ (final concentration of 2.5 mM) and placed on ice for 5 min.

A dilution series ranging from 0 to 30 µM of the respective recombinant PKM2 protein was prepared in PCR tubes using EMSA protein buffer (see Buffers, supplemental material; total volume: 10 µL per sample). Three independent protein preparations of PKM2 were used as biological replicates. To each sample, 1 µL of EMSA binding reaction buffer (see Buffers, supplemental material) and 10 µL of the prepared Cy5-labelled RNA were added (final RNA concentration: 10 nM; final recombinant protein concentration: 0 to 15 µM). Samples were incubated at 25 °C for 20 min in a thermocycler.

After pre-running a BioRad 4-15% Criterion™ TGX™ Precast Midi Protein gel (BioRad) for 15 - 20 min at 100 V and 4 °C in 1x native gel running buffer (see Buffers, supplemental material), 19.8 µL of the prepared samples were loaded per well and the gel was run protected from light for approximately 45 min at 100 V and 4 °C. Fluorescence signals were detected on a Typhoon™ laser scanner using 600 V and 50 µm pixel size settings on Cy5-channel. Fluorescence signals were quantified using the BioRad Image Lab software. All results shown are based on the background-corrected, adjusted volume and were normalized to the no protein control.

### Quantification and Statistical Analysis

Statistical analysis was performed by unpaired or paired Student’s t test or two- or one-way ANOVA without correction for multiple comparison (Fisher LSD test). Please see figure legends for detailed information. Significance levels were set at p* < 0.05, p** < 0.01 and p*** < 0.001. For statistical analysis, GraphPad Prism was used.

## RESULTS

### Replacement of RNase treatment by sonication-based RNA fragmentation reduces background

In contrast to many canonical RBPs bearing high affinity RBDs, non-canonical RBPs may associate with RNA only transiently or through a minor fraction of the total protein pool, increasing input requirements and rendering these proteins particularly sensitive to background RNA carry-over during IP.

Aiming to identify the RNAs that specifically interact with the glycolytic enzyme RBP pyruvate kinase M2 (PKM2), we first established HeLa cells as a suitable system for PKM2 CLIP-seq analysis. PKM2 RNA and protein were readily detectable under standard culture conditions (data not shown). To confirm RNA-binding activity by PKM2, we performed silica-column-based complex capture (2C) and radioactive PNK analyses. Both assays revealed crosslinking- and RNase-sensitive signals, indicative of RNA binding by PKM2 in HeLa cells (Supplementary Figure S1A, B). We further validated PKM2-associated RNA signals using a pCp assay, in which biotin-labeled pCp is ligated to RNA fragments using T4 RNA ligase after RNase digestion and protein IP, a protocol highly similar to CLIP-seq approaches (Figure 1A). Signal specificity was validated by PKM2 knockdown controls (Supplementary Figure S1C-E).

**Figure 1:**
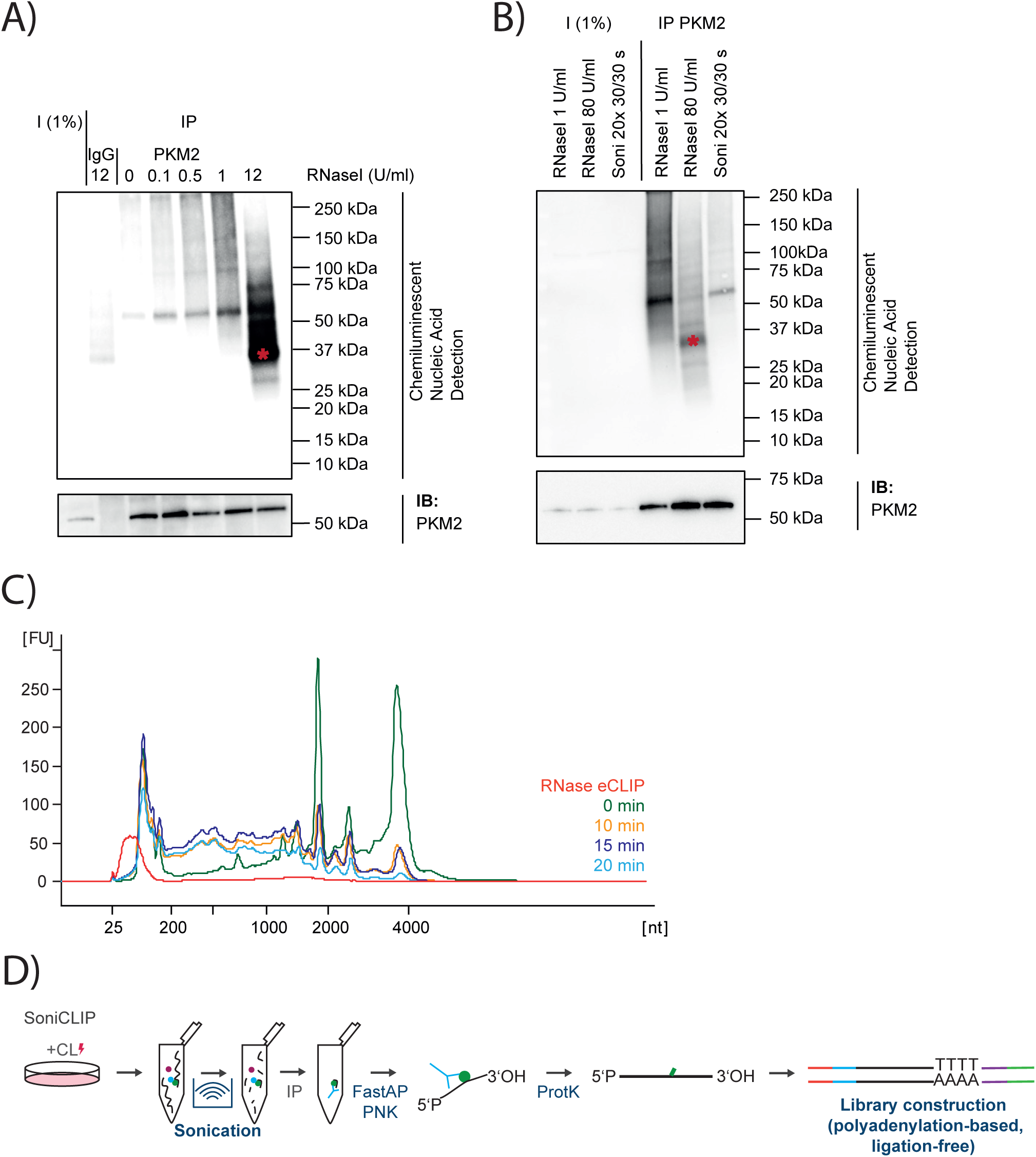
Replacement of RNase treatment by sonication-based RNA fragmentation reduces background. (A) pCp assay of PKM2 in HeLa cells. PKM2 was immunoprecipitated after crosslinking RNA-RBP complexes by UV-C light, cell lysis and RNA fragmentation. RNA was ligated to pCp-biotin and chemiluminescent nucleic acid detection was performed. Different concentrations of RNaseI were used. Immunoblot analysis of PKM2 from the same experiment. (*) background signal detected in samples with high RNaseI concentration. All samples were crosslinked. N=3. Representative blot is shown. (B) pCp assay of PKM2 in HeLa cells. RNase I concentration of 1 U/ml and 80 U/ml (standard eCLIP concentration) as well as sonication (20 cycles, 30 s on / 30 s off) are shown. Immunoblot analysis of PKM2 from the same experiment. (*) background signal detected in samples with high RNaseI concentration. All samples were crosslinked. N=2. Representative blot is shown. (C) RNA Bioanalyzer profiles of total RNA extracted from HeLa cell lysates after RNase I digestion or sonication. N=2. Representative experiment is shown. FU: fluorescent units. (D) Schematic summary of soniCLIP workflow.

During RNase titration in the pCp assay, we observed that background noise increased with RNase I concentration. At higher RNase I concentrations, the PKM2-associated RNA signal was masked by an intensive smear between 25 and 75 kDa, with a strong band detected below 37 kDa, which was also detected in the IgG control (Figure 1A). Given its apparent molecular mass, this signal is likely derived from recombinant RNase I itself, which has a calculated molecular mass of approximately 27 kDa and may be co-immunoprecipitated when present in excess despite stringent washing. Since this band represents labelled RNA fragments not covalently crosslinked to PKM2 *in vivo*, its carry-over could introduce substantial background into downstream CLIP-seq libraries despite stringent washing conditions. This is particularly relevant for gel-free CLIP approaches, in which all immunoprecipitated RNA fragments can enter library preparation. Notably, the background also extended into higher molecular weight regions that overlap with the expected RBP-RNA complexes (Figure 1A). These observations suggested that RNase-dependent fragmentation can introduce non-specific RNA, especially when profiling non-canonical RBPs with lower quantities of specifically bound RNAs.

To overcome this limitation, we explored sonication as an RNase-independent strategy for fragmenting crosslinked RNA. Sonication is routinely used to fragment crosslinked chromatin in ChIP-seq workflows (55,56), and we reasoned that it could similarly fragment RNA while avoiding the introduction of an RNA-bound enzyme into the IP reaction. Applying 20 cycles of sonication in the PKM2 pCp assay produced a distinct PKM2-associated RNA signal of appropriate mass at ∼60 kDa (Figure 1B). In contrast to RNase I-treated samples, sonicated samples showed strongly reduced background and lacked both the RNase-sized band and associated smears observed under RNase I digestion conditions representative of conventional CLIP-seq workflows (80 U/ml). These results indicate that sonication preserves crosslinked PKM2-RNA complexes while reducing non-specific RNA carry-over.

We next assessed whether sonication-generated RNA fragment lengths are compatible with CLIP-seq library preparation. Increasing the duration of sonication progressively shifted RNA fragments into the range of approximately 50-200 nucleotides (nt), comparable to fragment sizes typically obtained by RNase treatment for CLIP-seq analyses (Figure 1C). Thus, sonication provides an RNase-independent fragmentation strategy that yields sequencing-compatible RNA fragments while minimizing RNase-associated background.

Based on these findings, we systematically developed soniCLIP as a streamlined CLIP-seq workflow designed for low-input, gel-free and ligation-free profiling of RBPs (Figure 1D; Supplementary Figure S1F). In soniCLIP, UV crosslinking is followed by cell lysis, sonication-mediated RNA fragmentation and IP of the RBP-RNA complex of interest. After elution, protein-bound RNA fragments undergo 3′- and 5′-end repair, followed by proteinase K digestion to release RNA fragments. Libraries are then generated using the SMARTer smRNA-Seq Kit (Takara), which captures small RNA fragments through 3′ polyadenylation and ligation-free SMART-based cDNA synthesis. By combining RNase-independent fragmentation with ligation-free library preparation, soniCLIP eliminates gel purification and RNA ligation steps while retaining region-level identification of RBP-associated RNA fragments suitable for low-input analyses of canonical and non-canonical RBPs.

### soniCLIP supports reproducible library generation and robust read processing

To benchmark soniCLIP against an established CLIP framework, we selected two canonical RBPs with well-characterized binding properties and extensive public reference datasets: stem-loop binding protein (SLBP) and RNA-binding Fox-1 homolog 2 (RBFOX2). SLBP binds the conserved stem-loop element in the 3′ region of replication-dependent histone mRNAs and thus represents an RBP with a small, well-defined target set (21). RBFOX2, in contrast, recognizes the conserved 5’-UGCAUG-3’ sequence motif and binds more broadly, predominantly in intronic regions linked to splicing regulation (22). Together, the two RBPs provide complementary reference systems for assessing target recovery, motif specificity and reproducibility. Since both RBPs have been profiled by ENCODE eCLIP in K562 cells (4), we used this cell line to enable direct comparison with eCLIP, a widely used CLIP implementation.

We first optimized sonication-mediated RNA fragmentation in K562 cells. Sonication for 15-20 min efficiently fragmented both rRNA and mRNA, generating RNA fragments in a size range suitable for CLIP-seq library preparation (Figure 2A). Based on these results, we used 15 min of sonication for subsequent SLBP and RBFOX2 soniCLIP experiments. IP conditions were established for both RBPs using the polyclonal antibodies reported in the corresponding ENCODE eCLIP experiments (Supplementary Figure S2A and B) (4). For soniCLIP, 500 µg of total protein were used per IP, corresponding to 10% of the 5 mg input commonly used in ENCODE-style eCLIP workflows.

**Figure 2:**
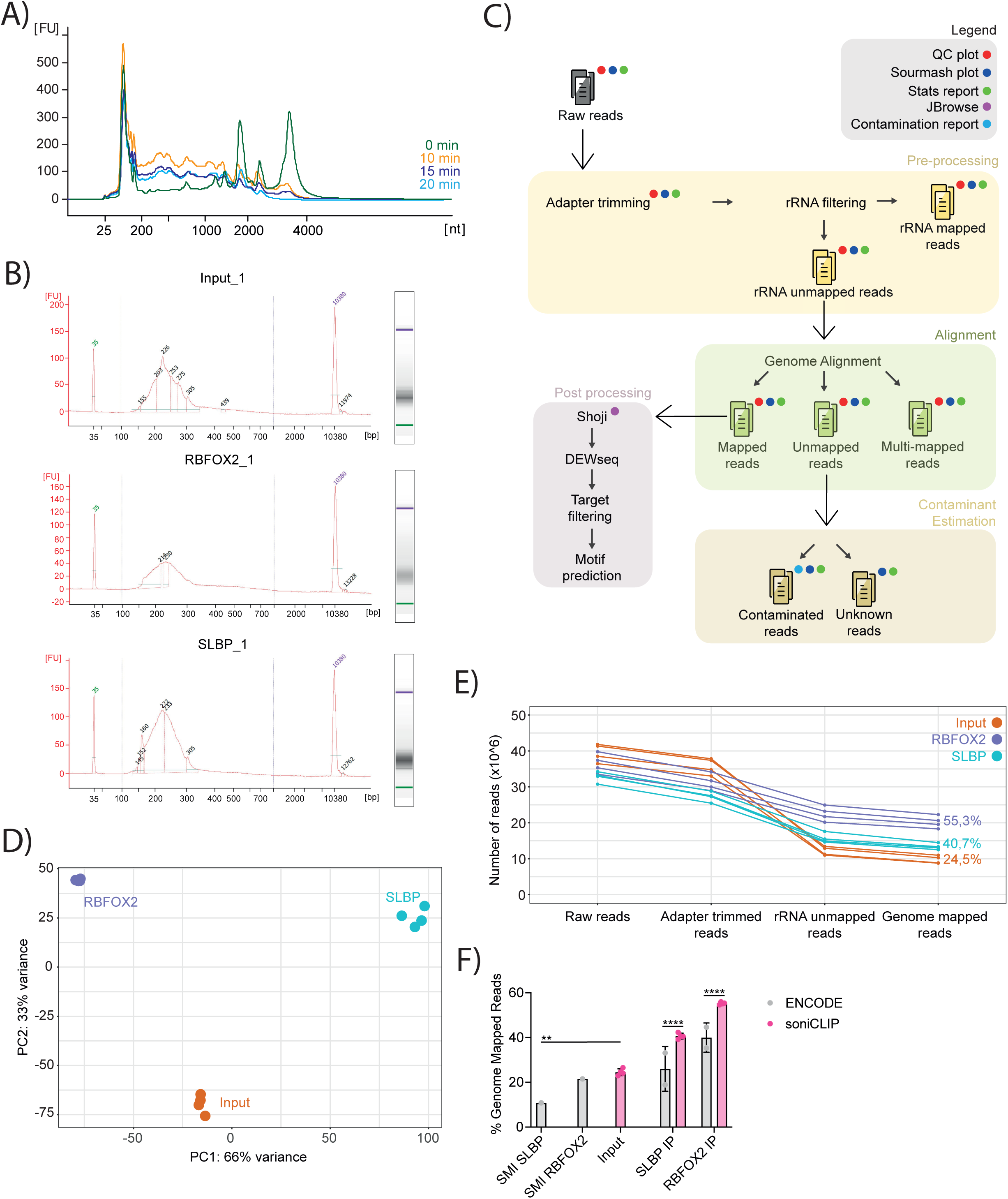
soniCLIP supports reproducible library generation and robust read processing. (A) RNA Bioanalyzer profiles of total RNA extracted from K562 cell lysates after sonication. N=2. Representative experiment is shown. (B) DNA Bioanalyzer profiles of soniCLIP libraries at the post-size selection stage. N=4. Representative libraries are shown. FU: fluorescent units. (C) soniCLIP analysis workflow. (D) PCA of the soniCLIP benchmarking dataset. (E) Read processing and alignment statistics of the soniCLIP benchmarking dataset. (F) Quantification of genome mapped reads (%) relative to raw reads for soniCLIP and ENCODE eCLIP data(4). Mean +/-SD. Two-way ANOVA. *p < 0.05; **p < 0.01; ***p < 0.001; ****p < 0.0001. SMI: size-matched input.

soniCLIP was then performed in biological quadruplicates for SLBP, RBFOX2 and input samples. Successful library generation was confirmed before and after size selection, which enriched for inserts below 150 nt (Figure 2B; Supplementary Figure S2C). All IP and input libraries were sequenced as a single multiplexed pool (Supplementary Table S1). We then established a streamlined soniCLIP data analysis workflow for rapid processing and target identification (Figure 2C) (see Methods section for details). Post adapter trimming, the reads were aligned to rRNA reference sequences. The rRNA-mapped reads were discarded and the rRNA-free reads were subsequently aligned to the human genome. The genome unmapped reads were assessed for potential library contamination, with no major contaminants being detected (data not shown). Genome mapped reads were processed using the shoji suite to extract crosslink sites and export them as a count matrix, along with a window-level annotation table and a max-count table. The aforementioned tables were then used for downstream target identification using DEWSeq (57). Briefly, reads were evaluated by applying sliding windows to detect local enrichment, and significant adjacent windows were subsequently merged into regions representing robust binding events. To enable direct comparison with ENCODE eCLIP data, publicly available ENCODE eCLIP datasets were re-aligned and processed using the same analysis workflow.

Several analysis steps were adapted to the specific properties of soniCLIP libraries. First, because soniCLIP is ligation-free, adapter re-ligation and double-ligation artifacts are avoided, enabling a straightforward one-step adapter trimming procedure, compared to the two-step adapter trimming in eCLIP. Second, as soniCLIP libraries arise from read-through rather than truncation at the crosslink site, the middle-site of the read was used as an approximate positional anchor for metaprofile analyses rather than as direct crosslink-site calls. Third, because CLIP datasets are sparse at the window level, several windows contain no reads in one or more samples, either because the region is not bound, the signal is close to the detection limit, or enrichment is present in the IP but absent from input or control samples. To account for this excess of zero values before differential enrichment testing, we incorporated ZINB-WaVE (48), which models zero-inflated count distributions and improves variance estimation. Finally, because soniCLIP does not include gel-based excision of a defined RBP-RNA complex, enriched regions were identified using both input and an unrelated IP as specificity controls. Thus, significant SLBP windows were defined as windows enriched over both input and RBFOX2 IP samples, and RBFOX2 windows were called analogously using input and SLBP IP samples as controls. For ENCODE eCLIP datasets, adapter trimming and crosslink-site assignment were performed according to protocol specifications described in (4) and enriched-region calling was performed using DEWseq (IP vs SMI). To enable a balanced comparison, soniCLIP and ENCODE eCLIP datasets were processed in parallel within the same general analysis framework, while adapting method-specific steps to the properties of each library type. In this way, both datasets were analyzed as consistently as possible without ignoring differences inherent to the respective experimental workflows.

Quality control analyses supported the reproducibility of soniCLIP. Hierarchical clustering of the sample level signatures showed separation of the biological replicates by sample group after trimming, and this separation increased after genome mapping (Supplementary Figure S2D). Principal component analysis further showed robust clustering of biological replicates (Figure 2D). Read lengths ranged from approximately 15 to 115 nt, with most reads being between 40 and 90 nt (Supplementary Figure S2E), a size distribution suitable for efficient genome alignment.

We next compared read retention during soniCLIP processing with published eCLIP data. In many CLIP-seq approaches, including eCLIP, substantial read loss occurs during preprocessing, with only a fraction of reads ultimately aligning to the genome (2). In soniCLIP, more than 95% of reads were retained after adapter trimming, consistent with the absence of ligation-associated adapter artifacts (Figure 2E). As expected, removal of rRNA-derived reads led to a subsequent decrease in read numbers. This decrease accounted for approximately 30-40% of reads in IP samples and was higher in input samples, where rRNA represents a larger fraction as the input sample is derived from total cellular RNA. After rRNA removal and genome alignment, an average of 24.5% of input reads, 40.7% of SLBP IP reads and 55.3% of RBFOX2 IP reads mapped to the genome (Figure 2E and F). In this benchmark analysis, soniCLIP yielded a higher fraction of genome-mapped reads than the corresponding ENCODE eCLIP SMI and IP datasets, supporting efficient read processing in the soniCLIP workflow (Figure 2F).

Together, these results show that soniCLIP supports reproducible library generation, efficient read processing and robust biological replicate clustering from low-input material. The marked reduction in read loss during processing highlights the advantage of ligation-free library preparation and provides a strong basis for downstream identification of RBP-associated RNA regions.

### Identification of SLBP targets by soniCLIP from low-input material

We next assessed soniCLIP performance in greater detail using SLBP, a canonical RBP with a well-defined target RNA repertoire (21). For best comparability, soniCLIP and ENCODE eCLIP datasets were processed with the same DEWSeq-based analysis workflow (Supplementary Table S2, S3; see Methods).

For soniCLIP, enriched SLBP windows were required to be significant both over input and over the unrelated RBFOX2 IP sample (Supplementary Figure S3A). This two-control strategy accounts for transcript abundance as well as IP-related background that may be shared across unrelated pulldowns. ENCODE eCLIP SLBP windows were identified according to the eCLIP strategy by comparison with the corresponding SMI control (Supplementary Figure S3B). This analysis provided the basis for region-level comparison of SLBP target recovery between soniCLIP and eCLIP.

We first analyzed significant regions using stringent thresholds for soniCLIP (padj ≤0.01, Log_2_FC ≥2), which identified 114 SLBP-enriched regions (Figure 3A and B). Applying the same thresholds to the ENCODE eCLIP dataset yielded 35 regions, likely reflecting that ENCODE eCLIP only uses two IP replicates and one SMI control compared with the 4 replicates per experimental condition used for soniCLIP. We therefore additionally evaluated eCLIP with a more permissive threshold (padj ≤0.1, Log_2_FC ≥1), which identified 150 enriched regions. For benchmarking, we compare soniCLIP results to permissively analyzed ENCODE eCLIP to allow for a more balanced comparison, while retaining the stringent eCLIP results for reference.

**Figure 3:**
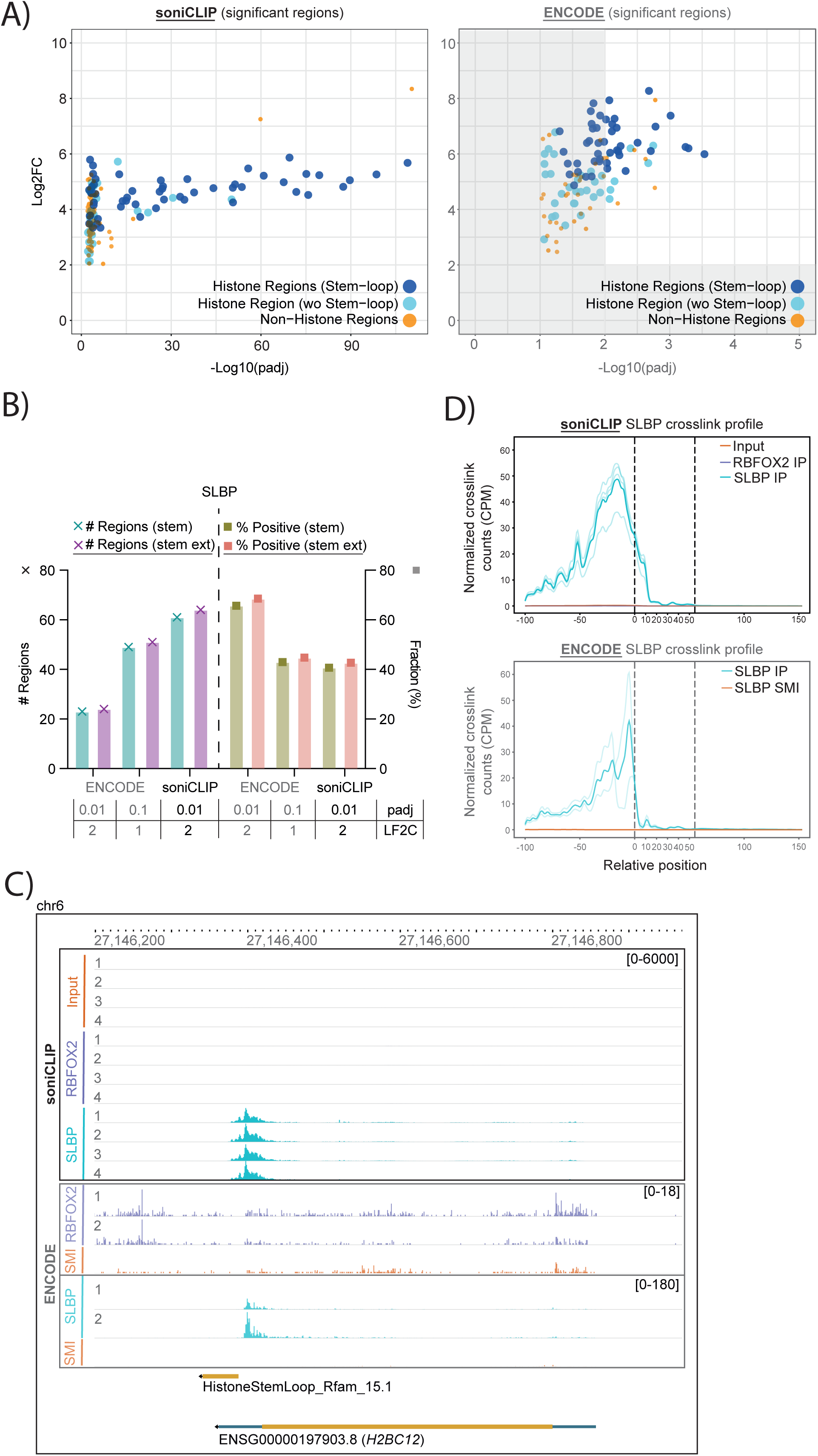
Identification of SLBP targets by soniCLIP recovers SLBP-bound histone stem-loop targets from low-input material. (A) Significant regions in SLBP soniCLIP (padj ≤0.01, L2FC ≥2) and SLBP ENCODE eCLIP analysis (grey scale: padj ≤0.01, L2FC ≥2; padj ≤0.1, L2FC ≥1). Histone-regions with and without stem-loops are highlighted. (B) Quantification of significant histone stem-loop containing regions (X) and fractions (◻) in soniCLIP and ENCODE eCLIP SLBP data. Presence of conserved histone stem-loop regions were analyzed with more stringent (stem) and more relaxed (stem ext) settings (see methods). (C) Exemplary genome browser tracks showing projected crosslink sites for soniCLIP and ENCODE eCLIP data. *H2BC12* is shown. (D) SLBP crosslink profiles for soniCLIP and ENOCDE eCLIP are shown. Crosslink profiles are plotted using CPM (Counts Per Million) normalized crosslink counts for ± 100 nucleotide window around histone 3’ UTR stem loop reference locations.

Using histone stem-loop structures (stem) as the expected SLBP target feature, soniCLIP identified 61 stem-loop-containing regions, compared with 49 regions in the permissive ENCODE eCLIP analysis (Figure 3B). Thus, despite using only 10% of the starting material, soniCLIP recovered additional histone stem-loop-containing regions, corresponding to a 25% increase in target discovery. The proportion of enriched regions containing a conserved histone stem-loop was comparable between soniCLIP and eCLIP (41% and 43%, respectively), indicating that the increased recovery did not come at the expense of target specificity. When less stringent settings were used for stem-loop annotation (stem ext), both datasets showed a modest further increase in recovery, with three additional regions detected by soniCLIP and two additional regions detected by eCLIP (Figure 3B).

Analysis of representative SLBP binding sites further supported robust soniCLIP performance. soniCLIP signals were highly reproducible across biological replicates and showed clear enrichment over input and unrelated IP controls, facilitating target identification and contributing to the increased recovery of histone stem-loop-containing regions (Figure 3C; Supplementary Figure S3C). We next assessed the distribution of projected crosslink sites around histone stem-loop structures. Since soniCLIP libraries are generated with read-through across the crosslink sites, rather than reverse-transcription truncation as in eCLIP, crosslink positions were approximated at the middle-site of each soniCLIP read. Still, projected soniCLIP crosslink profiles recapitulated the relative positioning observed in ENCODE eCLIP data around histone stem-loop structures (Figure 3D). Moreover, crosslink profiles were more consistent between biological replicates in soniCLIP than in the corresponding eCLIP dataset.

Together, these results show that soniCLIP robustly recovers established SLBP target regions with specificity comparable to the ENCODE eCLIP benchmark, while requiring substantially less starting material. The recovery of additional histone stem-loop-containing regions, combined with reproducible replicate behavior and comparable projected crosslink profiles, supports the use of soniCLIP for robust low-input identification of structured RBP target sites.

### Identification of RBFOX2 motif-containing RNA regions with high specificity by soniCLIP from low-input material

We also assessed soniCLIP performance using RBFOX2, a canonical RBP with a broader target repertoire than SLBP. RBFOX2 regulates alternative splicing and predominantly binds to the conserved 5’-UGCAUG-3’ motif, which occurs far more frequently within the transcriptome than the histone stem-loop element recognized by SLBP (22). RBFOX2 therefore provides a complementary benchmark to evaluate soniCLIP performance on a canonical RBP with motif-specific but more widespread RNA binding.

As for SLBP, soniCLIP and ENCODE eCLIP datasets were analyzed using the same DEWSeq-based workflow (Supplementary Table S4, S5). For soniCLIP, significant RBFOX2 windows were defined as the overlap between windows enriched over input and windows enriched over the unrelated SLBP IP sample, thereby controlling for both transcript abundance and IP-related background (Supplementary Figure S4A). ENCODE eCLIP RBFOX2 enrichment was assessed against the corresponding SMI control (Supplementary Figure S4B). Region-level analysis was then performed using the same thresholding strategy as for SLBP. Since stringent cutoffs identified only 20 significant regions in the ENCODE eCLIP dataset, the relaxed eCLIP threshold was used for the main comparison, with the results applying the stringent cutoff parameters shown for reference.

soniCLIP identified 74 significant RBFOX2 regions containing the 5’-UGCAUG-3’ motif, corresponding to 40% of all significant regions (Figure 4A, B). ENCODE eCLIP recovered a higher absolute number of motif-containing regions (106), but with a lower motif-containing fraction (36%). Inspection of representative binding sites further showed strong reproducibility between biological replicates in the soniCLIP datasets (Figure 4C; Supplementary Figure S4C).

**Figure 4:**
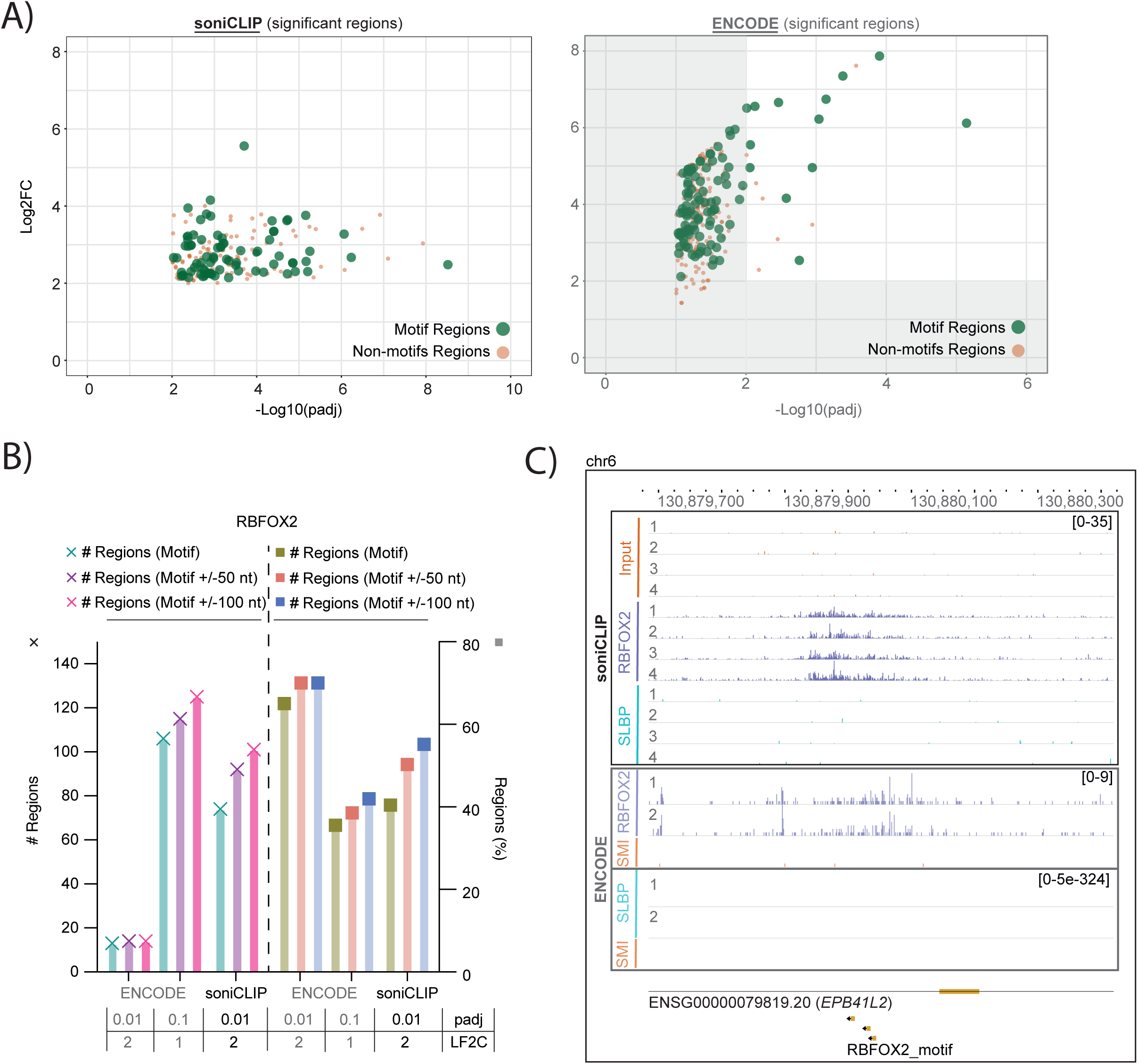
Identification of RBFOX2 motif-containing RNA regions with high specificity by soniCLIP from low-input material. (A) Significant regions in RBFOX2 soniCLIP (padj ≤0.01, L2FC ≥2) and RBFOX2 ENCODE eCLIP analysis (grey scale: padj ≤0.01, L2FC ≥2; padj ≤0.1, L2FC ≥1). Regions with and without RBFOX2 motif are highlighted. (B) Quantification of significant motif containing regions (X) and fractions (◻) in soniCLIP and ENCODE eCLIP RBFOX2 data. Presence of the RBFOX2 motif within the significant region or in a window including the significant region and an area ±50 nt or ±100 nt up-/downstream of the significant region was analyzed (see methods). (C) Exemplary genome browser tracks showing projected crosslink sites for soniCLIP and ENCODE eCLIP data. *EPB41L2* is shown.

Since soniCLIP is not based on reverse-transcription truncation at the crosslink site and, importantly, because the crosslink site and the biological binding site are not necessarily identical (57), we also assessed motif recovery in areas adjacent to significant regions (±50 or ±100 nt, respectively). This increased both the number and the fraction of 5’-UGCAUG-3’-containing regions in both datasets. When including motifs ±100 nt up-/downstream of significant regions, soniCLIP identified 101 motif-containing regions, corresponding to 55% of significant regions, whereas ENCODE eCLIP identified 125 motif-containing regions, corresponding to 42% (Figure 4B). The stronger effect on soniCLIP is consistent with the read-through nature of the library preparation, in which enriched fragments can extend into motif-adjacent regions rather than marking the crosslink site at nucleotide resolution.

Taken together, the RBFOX2 analyses show that soniCLIP recovers motif-containing RBFOX2-associated regions with high reproducibility and motif specificity comparable to the ENCODE eCLIP benchmark. While eCLIP recovered a higher absolute number of 5’-UGCAUG-3’-containing regions, soniCLIP achieved a higher motif-containing fraction and robust replicate reproducibility from only 10% of the typical starting material. Combined with the SLBP analysis, these results demonstrate that soniCLIP recovers both highly restricted structured targets and broader motif-defined binding landscapes with sensitivity and specificity comparable to ENCODE eCLIP benchmarks.

### Robust target identification for the non-canonical RBP PKM2 by soniCLIP

Based on the above results, we used soniCLIP for profiling a non-canonical RBP. CLIP-based approaches often yield suboptimal data for this class of RBPs, with reduced signal-to-noise ratios, increased read loss during processing, and limited reproducibility between biological replicates (3). These challenges likely reflect the transient nature of many non-canonical RNA-protein interactions and the frequently limited fraction of the protein of interest that is RNA-bound at steady state. We therefore applied soniCLIP to PKM2 in HeLa cells; to our knowledge, no HeLa-derived eCLIP data have been published for PKM2. Sonication conditions had been established previously (Figure 1C), and PKM2 IP was further optimized before library preparation (Supplementary Figure S5A).

soniCLIP libraries were generated as described above (Figure 1D; Supplementary Figure S1F; Supplementary Table S6), and library quality was assessed before and after size selection (Figure 5A; Supplementary Figure S5B). As for the canonical RBPs profiled in this study, IPs were performed from only 500 µg total protein starting material. In parallel, SLBP was included as an unrelated IP control, enabling assessment of both input- and IP-associated background by overlapping DEWSeq analysis. Libraries were sequenced as a multiplexed pool and processed using the established soniCLIP analysis pipeline with minor modifications (Figure 2C). Since PKM2 has previously been reported to interact with rRNA, reads were mapped to a reference genome supplemented with a synthetic, *in silico* ribosomal chromosome, allowing ribosomal targets to be retained and evaluated during analysis (see Methods section for details).

**Figure 5:**
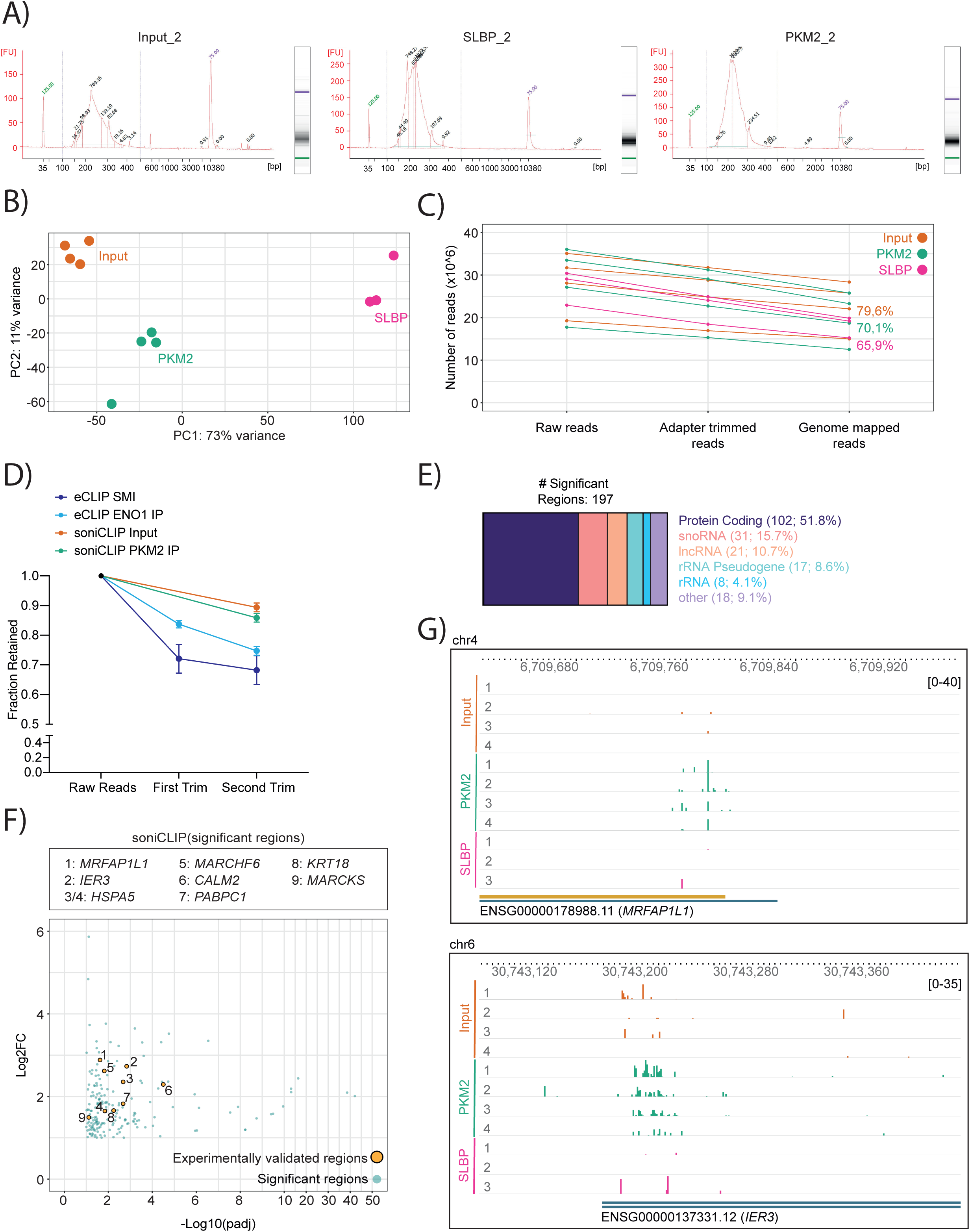
Robust target identification for the non-canonical RBP PKM2 by soniCLIP. (A) DNA Bioanalyzer profiles of soniCLIP libraries at the post-size selection stage. Representative libraries are shown. N=4. (B) PCA of the soniCLIP PKM2-SLBP dataset. (C) Read processing and alignment statistics of the soniCLIP PKM2-SLBP dataset. (D) Read processing statistics of the soniCLIP PKM2-SLBP dataset in comparison to ENO1 eCLIP data (58). (E) Gene type assignment of the 197 significantly enriched PKM2 regions. (F) Significant regions in the PKM2 soniCLIP dataset (padj ≤0.1, Log_2_FC ≥1). Experimentally validated regions are highlighted. (G) Exemplary genome browser tracks showing projected crosslink sites for PKM2-SLBP soniCLIP data. *MRFAP1L1* and *IER3* are shown.

Quality-control analyses indicated robust soniCLIP performance for PKM2. Hierarchical clustering of the sample level signatures and principal component analysis showed separation of the biological replicates, demonstrating a level of reproducibility that is often difficult to achieve when profiling non-canonical RBPs by other CLIP-based methods (Figure 5B; Supplementary Figure S5C). Read length distributions were within the expected range, with most reads being between 40 and 90 nts (Supplementary Figure S5D), and more than 65% of reads mapped to the genome (Supplementary Figure S5C). The high mapping rate was supported by inclusion of the synthetic ribosomal chromosome in the adapted reference genome. Compared with eCLIP datasets previously generated in our laboratory for non-canonical RBPs (58), soniCLIP retained a markedly higher fraction (>10%) of reads after adapter trimming (Figure 5D). This finding supports the rationale that ligation-free library preparation reduces read loss, particularly in low-signal applications such as non-canonical RBP profiling. Together, these results show that soniCLIP generates high-quality, reproducible libraries from low-input PKM2 IPs.

We then used the PKM2 soniCLIP dataset to identify candidate RNA targets. Differential enrichment was assessed by DEWSeq analysis using padj ≤0.1 and Log_2_FC ≥1 (Supplementary Table S7). These relatively permissive cutoffs were chosen to support broader *de novo* discovery, as is commonly useful for non-canonical RBPs where RNA-binding properties, target spectra and expected enrichment patterns are less well established. As for the canonical RBPs, significant regions were defined by overlapping windows enriched in PKM2 IP compared with both the unrelated SLBP IP and input controls (Supplementary Figure S5E). This analysis identified 197 candidate PKM2-associated RNA regions (Figure 5E, F). Motif enrichment analysis did not reveal a dominant sequence motif, consistent with the possibility that PKM2 recognizes structural or context-dependent RNA features rather than a short linear sequence motif. In line with previous reports (25), a subset of candidate regions mapped to rRNA (25 regions; 12.7%) with enrichment among regions aligning to the 18S rRNA reference (Supplementary Table S7). The majority of candidate regions, however, mapped to protein-coding genes (102 regions, 51.8%). Several significantly enriched regions were located in untranslated regions (UTRs), including candidate PKM2-binding sites in the 5′-UTR of *MRFAP1L1* and the 3′-UTR of *IER3* (Figure 5G). These results demonstrate that soniCLIP can reproducibly identify candidate RNA targets of a non-canonical RBP, detecting both ribosomal and mRNA-associated interactions.

### Experimental validation of PKM2 RNA targets identified by soniCLIP

To evaluate the validity of the candidate PKM2 targets identified by soniCLIP, we first performed RIP-qRT-PCR for 12 candidate targets using TaqMan-based detection chemistry (Figure 5F; Figure 6A). For each target, probes were selected within regions identified as significantly enriched by soniCLIP, thereby directly testing the candidate binding regions detected by DEWSeq analysis. Eight of the twelve tested targets (67%) were significantly enriched over background.

**Figure 6:**
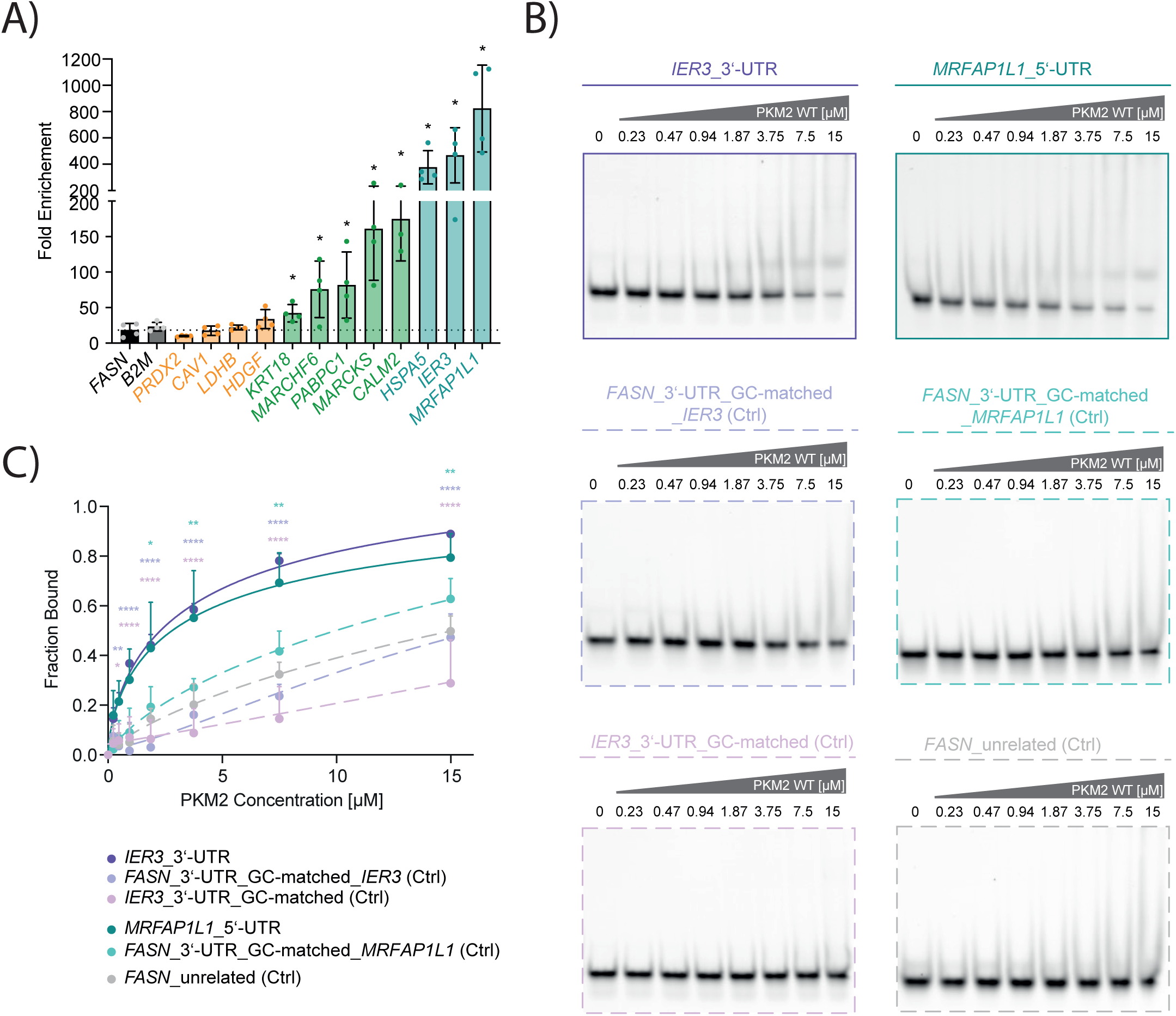
Experimental validation of PKM2 RNA targets identified by soniCLIP. (A) RIP-qRT-PCR analysis of cDNA derived from *in cellulo* PKM2-crosslinked RNA. *FASN* and *B2M* were used to assess background enrichment over IgG control. Significance levels were calculated compared to the *FASN* control. Mean +/-SD. One-way ANOVA. *p < 0.05; **p < 0.01; ***p < 0.001; ****p < 0.0001. (B) EMSA analysis of recombinant WT PKM2 and Cy5-labelled RNA target or control oligos. N=3 (for each biological replicate an independent recombinant protein preparation was used). Representative EMSAs are shown. (C) EMSA signal quantification. RNA-bound fraction is shown. Mean +SD. Two-way ANOVA. *p < 0.05; **p < 0.01; ***p < 0.001; ****p < 0.0001.

To further assess direct binding, we performed electrophoretic mobility shift assays (EMSAs) with recombinant PKM2 and synthetic RNA oligonucleotides (Figure 6B, C). Oligonucleotides of 35-45 nts were designed from significantly enriched regions in the *IER3* 3′-UTR and the *MRFAP1L1* 5′-UTR. Size- and GC-content-matched control oligonucleotides were included to assess non-specific binding. Recombinant PKM2 bound specifically to both target-derived oligonucleotides compared with their respective controls, validating direct interaction with RNA sequences derived from soniCLIP enriched regions (Figure 6B, C).

Although soniCLIP does not preserve single-nucleotide crosslink-site information in the same way as ligation-based CLIP approaches, these results show that enriched regions are resolved with sufficient precision to guide experimental validation of previously uncharacterized RBP-RNA interactions. Together with the sequencing quality-control metrics, replicate reproducibility and reduced read loss described above, the validation experiments demonstrate that soniCLIP enables robust identification of RNA targets for the non-canonical RBP PKM2 in a low-input setting.

## DISCUSSION

CLIP-based technologies have transformed the analysis of RNA-protein interactions from living cells, but their implementation remains challenging. Most protocols require careful balancing of RNA fragmentation, stringent enrichment of crosslinked complexes, recovery of limited RNA material and construction of low-bias sequencing libraries (2). These requirements are manageable for high affinity RBPs and experienced laboratories, but they become limiting when the protein of interest has weak or transient RNA-binding activity. This is particularly relevant for non-canonical RBPs, which often lack classical RBDs and may associate with RNA only in specific molecular states or cellular contexts. The results presented here offer a practical, validated alternative for any CLIP experiment, and especially under challenging settings.

Replacing RNase digestion with sonication addresses a central source of variability in CLIP experiments. RNase treatment is widely used because it generates short RNA fragments suitable for binding-site mapping, but its outcome depends strongly on enzyme concentration, reaction conditions and sample composition (8–10). In addition, RNase itself can become a source of RNA-associated background (Figure 1A, B). While this observation was made in the PKM2 context, it highlights a general consideration for low-signal CLIP experiments: when the specific crosslinked RBP-RNA fraction is low, even modest background introduced during fragmentation or IP can dominate the signal. For non-canonical RBPs, reducing such background may be more critical than single nucleotide resolution. Sonication provides a physical alternative that avoids adding an RNA-binding nuclease to the sample and thereby reduces one route of nonspecific RNA carry-over.

Physical fragmentation also changes the balance of what a CLIP experiment is optimized to achieve. Classical CLIP approaches often aim to define crosslink sites at or near nucleotide resolution (3), whereas soniCLIP prioritizes reproducible identification of enriched RNA regions. This comes with an expected trade-off: soniCLIP does not preserve single-nucleotide crosslink-site information in the same way as methods that use reverse-transcription stops or mutation signatures (4–6,10). However, the value of nucleotide-level information depends on the quality and interpretability of the underlying signal. For low-signal RBPs, apparent crosslink-site resolution may be less useful if target recovery is poor, background is high or reproducibility between replicates is limited. It also needs to be considered that the UV-crosslink sites are not necessarily identical to the biological binding sites (57); in such cases, high resolution crosslink site identification can even be misleading. The validation of PKM2-enriched regions by RIP-qRT-PCR and EMSA shows that regional resolution is sufficient to guide downstream biochemical analysis. Thus, soniCLIP is best viewed as a target-discovery method on a region resolution level that emphasizes robustness and sensitivity over precise crosslink-site assignment.

The ligation-free design further supports this goal. Adapter ligation is a recurring bottleneck in eCLIP library preparation because it can introduce double- or re-ligated adapters that complicates downstream bioinformatic analysis and can lead to substantial read losses (2). These effects are particularly problematic in low-input settings, where each successive loss reduces the number of informative molecules available for analysis. By using a polyadenylation- and SMART-based library strategy, soniCLIP avoids direct RNA adapter ligation and reduces read loss during preprocessing. This is relevant not only for sequencing efficiency, but also for the statistical power of downstream enrichment analysis. A larger fraction of usable reads allows more robust comparisons between IP and control conditions, which is especially important for non-canonical RBPs with modest enrichment.

The performance of soniCLIP for the benchmark RBPs SLBP and RBFOX2 places the method in the context of established CLIP approaches. These proteins represent useful benchmark systems because their binding specificities are well documented and their expected target distributions are distinct. Our data show that soniCLIP identifies known RNA targets at least at par with eCLIP, a widely adopted method also applied to the large-scale ENCODE profiling (4,12,20) while using substantially less input. soniCLIP does not aim to replace existing CLIP variants, which remain preferable when nucleotide-resolution crosslink-site information is the main objective. Instead, it addresses a different need: reliable target recovery from lower input material and a streamlined experimental protocol.

This distinction is particularly important for expanding CLIP beyond specialized use cases. Many laboratories are interested in RNA-protein interactions but do not perform CLIP routinely, and protocol complexity can become a practical barrier. Gel purification, ligation optimization and extensive troubleshooting often require experience that is difficult to establish for occasional experiments. By removing gel-based purification and RNA adapter ligation, soniCLIP reduces the number of technically sensitive steps, makes the workflow easier to implement and reduces the hands-on time to only 3.5 days. This may also be valuable for rare primary cells and *in vivo* samples as well as candidate RBPs emerging from proteome-wide RNA-interactome studies, where material is limited and extensive method optimization is often not feasible.

The computational workflow contributes to the overall aims of soniCLIP. CLIP datasets are sparse and background-prone, and these features are amplified in low-input experiments or with non-canonical RBPs (59). The use of window-based analysis with DEWSeq, accounting for zero inflation through ZINB-WaVE and the consideration of enrichment over both input and unrelated IP controls provide a conservative framework for candidate region calling. This dual-control strategy is an important component of soniCLIP, as this gel-free approach does not generate a conventional size-matched input control like eCLIP (4). Computational background correction is therefore not an auxiliary step, but an integral part of the method. This is particularly relevant for gel-free workflows, where alternative control strategies are needed to account for nonspecific RNA recovery and IP-associated background (60). The input control accounts for transcript abundance and general RNA recovery, whereas the unrelated IP control helps to identify and remove antibody- or bead-associated RNA background, non-specifically bound abundant RNAs and regions enriched as a consequence of general IP behavior rather than RBP-specific association. Together, these normalizations help distinguish specific RBP-associated regions from general input signal and IP-associated contaminants, improving confidence in binding site detection. At the same time, the choice of unrelated IP control requires consideration. A control RBP with overlapping localization, abundance, RNA-binding mode or target class could remove genuine signal. Future soniCLIP experiments should therefore include input controls and, where possible, an unrelated IP control matched for sample type, while remaining biologically distinct from the RBP of interest.

Our PKM2 dataset illustrates the practical utility of soniCLIP for proteins whose RNA-binding mode is not yet well defined. PKM2 is a metabolic enzyme lacking canonical RBDs, and its interaction with RNA is therefore unlikely to follow the same rules as proteins that bear such RBDs (19). The absence of a dominant enriched sequence motif is consistent with the possibility that PKM2 recognizes RNA via structural or context-dependent features (24), although this will require further testing. Importantly, soniCLIP identified candidate regions that could be taken directly into successful validation assays, including short RNA fragments from the *IER3* 3′-UTR and *MRFAP1L1* 5′-UTR (Figure 6). This supports the use of soniCLIP as a discovery platform for non-canonical RBPs.

Several limitations should be considered when applying soniCLIP. Since fragmentation is achieved by sonication rather than RNase digestion, the method does not directly yield the same crosslink-site information obtained from some CLIP variants. Fragment boundaries should therefore not be interpreted as precise protein-contact sites. As with other antibody-based approaches, performance will depend on efficient and specific IP of the protein of interest. In addition, soniCLIP does not incorporate unique molecular identifiers (UMIs), which are used in some protocols to distinguish PCR duplicates from independently captured molecules (2). This represents a trade-off of the ligation-free, streamlined library design. However, soniCLIP is primarily intended for robust identification of enriched target regions rather than absolute molecule counting or nucleotide-level quantification. In this setting, the absence of UMIs represents an acceptable trade-off when library complexity is sufficient, PCR amplification is limited and candidate regions are defined by reproducible enrichment over both input and unrelated IP controls. Finally, while the reduced input requirement broadens applicability, very low-abundance proteins or interactions restricted to rare cellular states may still require further optimization of crosslinking, IP or sequencing depth.

Overall, soniCLIP should be particularly useful for laboratories seeking an accessible CLIP workflow, for studies with limited starting material and for the expanding field of non-canonical RBPs where robust target identification remains a major bottleneck.

## ACKNOWLEDGEMENTS

We thank current and former members of the Hentze laboratory for their feedback. We acknowledge EMBL’s core facilities, specifically the Genomics and the Protein Expression and Purification core facilities for their expert services, with special thanks to V. Benes, K. Remans and J. Scheurich.

## AUTHOR CONTRIBUTIONS

Conceptualization, P.S. and M.W.H.; Methodology, P.S., S.S., T.S., S.C., and D.F.-A.; Investigation, P.S., S.S., and T.S.; Writing - Original Draft, P.S. and M.W.H.; Writing - Review and Editing, P.S., S.S., T.S., S.C., D.F.-A, and M.W.H; Supervision, M.W.H.

## DATA AVAILABILITY

The soniCLIP benchmark and the PKM2 data have been deposited to GEO, under the accession numbers: GSE338605 and GSE338606, respectively.

Genome annotation and other reference data used in this analysis are available from the following DOI: 10.5281/zenodo.21334976. The data analysis workflow described above was developed using Nextflow, and has been submitted to WorfklowHub registry (https://workflowhub.eu/workflows/2197), and the computational analysis steps made use of the HPC resources provided by the EMBL IT services (https://doi.org/10.5281/zenodo.12785829). Shoji, for post-processing CLIP alignment data, is available under https://github.com/EMBL-Hentze-group/Shoji. Ngs-statter, a helper package for generating read statistics and certain plots is available under https://github.com/EMBL-Hentze-group/ngs-statter. DEWSeqHelper package is available under https://github.com/EMBL-Hentze-group/dewseq_helper.

## SUPPLEMENTARY DATA

Supplementary Table 1: Overview library preparation soniCLIP benchmark K562

Supplementary Table 2: DEWSeq analysis SLBP soniCLIP benchmark

Supplementary Table 3: DEWSeq analysis SLBP ENCODE eCLIP

Supplementary Table 4: DEWSeq analysis RBFOX2 soniCLIP benchmark

Supplementary Table 5: DEWSeq analysis RBFOX2 ENCODE eCLIP

Supplementary Table 6: Overview library preparation soniCLIP PKM2-SLBP HeLa

Supplementary Table 7: DEWSeq analysis PKM2 soniCLIP PKM2-SLBP

---

**Supplementary Figure 1:**
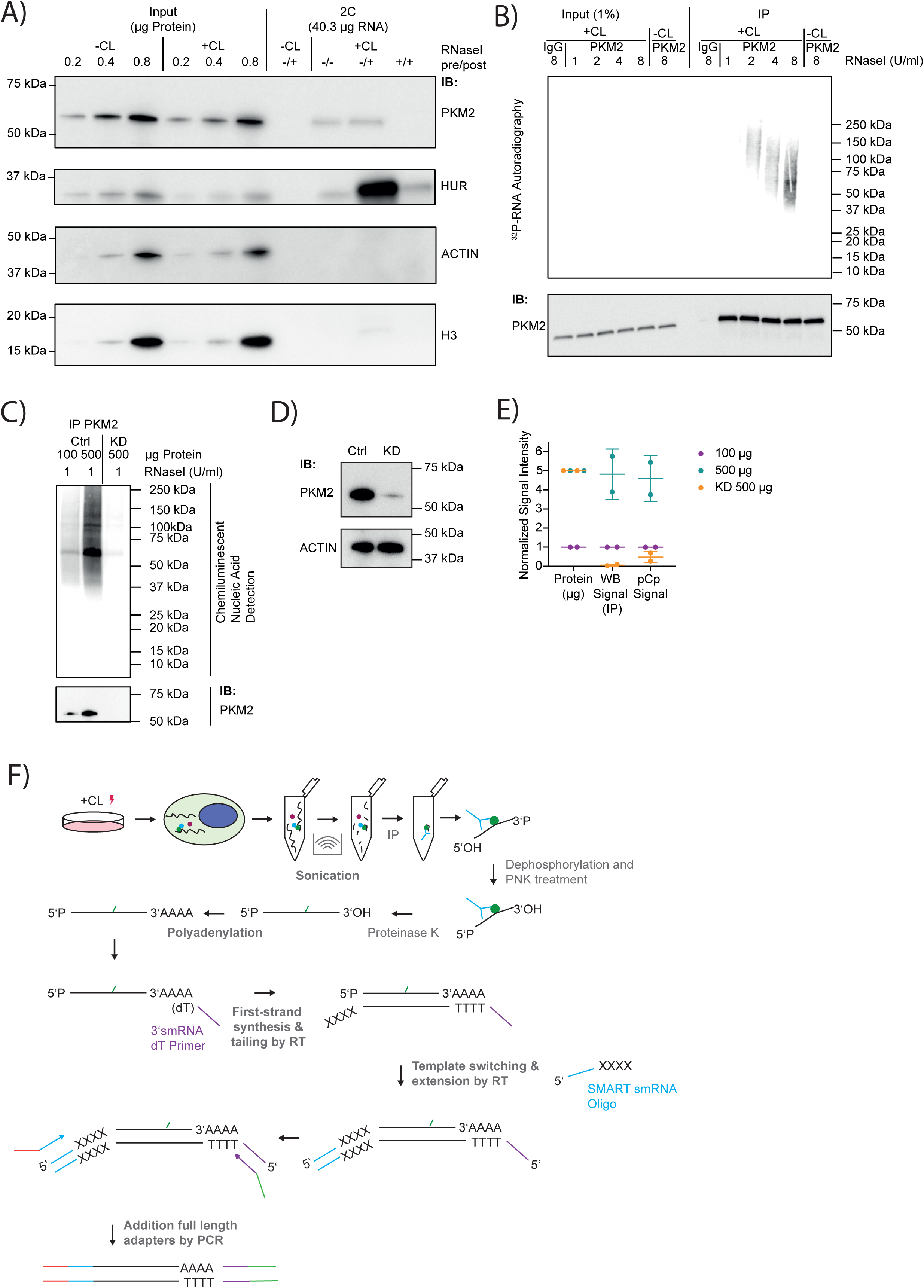
Replacement of RNase treatment by sonication-based RNA fragmentation reduces background. (A) Complex capture (2C) analysis of HeLa cell extracts. Immunoblots of PKM2, positive controls (HUR) and negative controls (ACTIN, H3) are shown. CL: crosslink. N=3. Representative blot is shown. (B) PNK assay of PKM2 in HeLa cells. PKM2 was immunoprecipitated after crosslinking RNA-RBP complexes by UV-C light, cell lysis and RNA fragmentation. Polynucleotide kinase labels RNA with radioactive 32P-yATP. Different concentrations of RNaseI were used. Immunoblot analysis of PKM2 from the same experiment. N=2. Representative blot is shown. (C) pCp assay of PKM2 in HeLa cells. PKM2 was immunoprecipitated after crosslinking RNA-RBP complexes by UV-C light, cell lysis and RNA fragmentation. RNA was ligated to pCp-biotin and chemiluminescent nucleic acid detection was performed. Western blot analysis of PKM2 from the same experiment. All samples were crosslinked. Ctrl: control siRNA, KD: knockdown. N=2. Representative blot is shown. (D) Western blot analysis of control and PKM2 KD HeLa cell extracts. PKM2 and ACTIN are shown. Ctrl: control siRNA, KD: knockdown. N=2. Representative blot is shown. (E) Quantification of pCp assay (S1C) and western blot (S1D) analysis of control and PKM2 KD HeLa cells. Ctrl: control siRNA, KD: knockdown. N=2. (F) Detailed soniCLIP workflow. Library construction is based on the SMARTer® smRNA-Seq Kit for Illumina® from Takara.

**Supplementary Figure 2:**
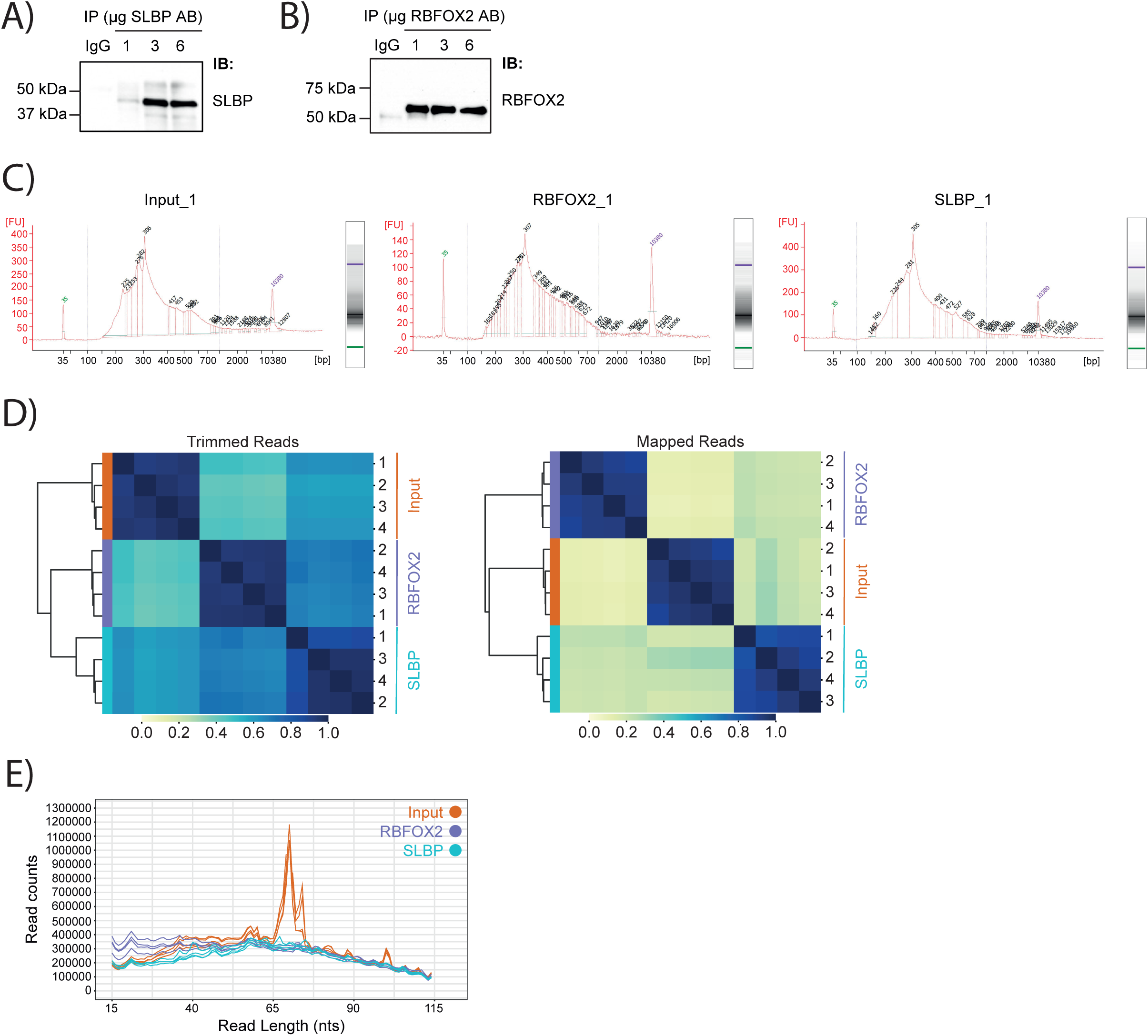
soniCLIP supports reproducible library generation and robust read processing. (A) Immunoblot of immunoprecipitated SLBP from K562 cell lysates. N=1. (B) Immunoblot of immunoprecipitated RBFOX2 from K562 cell lysates. N=1. (C) DNA Bioanalyzer profiles of soniCLIP libraries at the pre-size selection stage. Representative libraries are shown. N=4. (D) Sourmash plots of soniCLIP benchmark dataset trimmed reads and genome mapped reads as a means to visualize sequence similarity. Hierarchical clustering of similarity profiles scale 0-1. (E) Read length distribution soniCLIP benchmark dataset genome mapped reads.

**Supplementary Figure 3:**
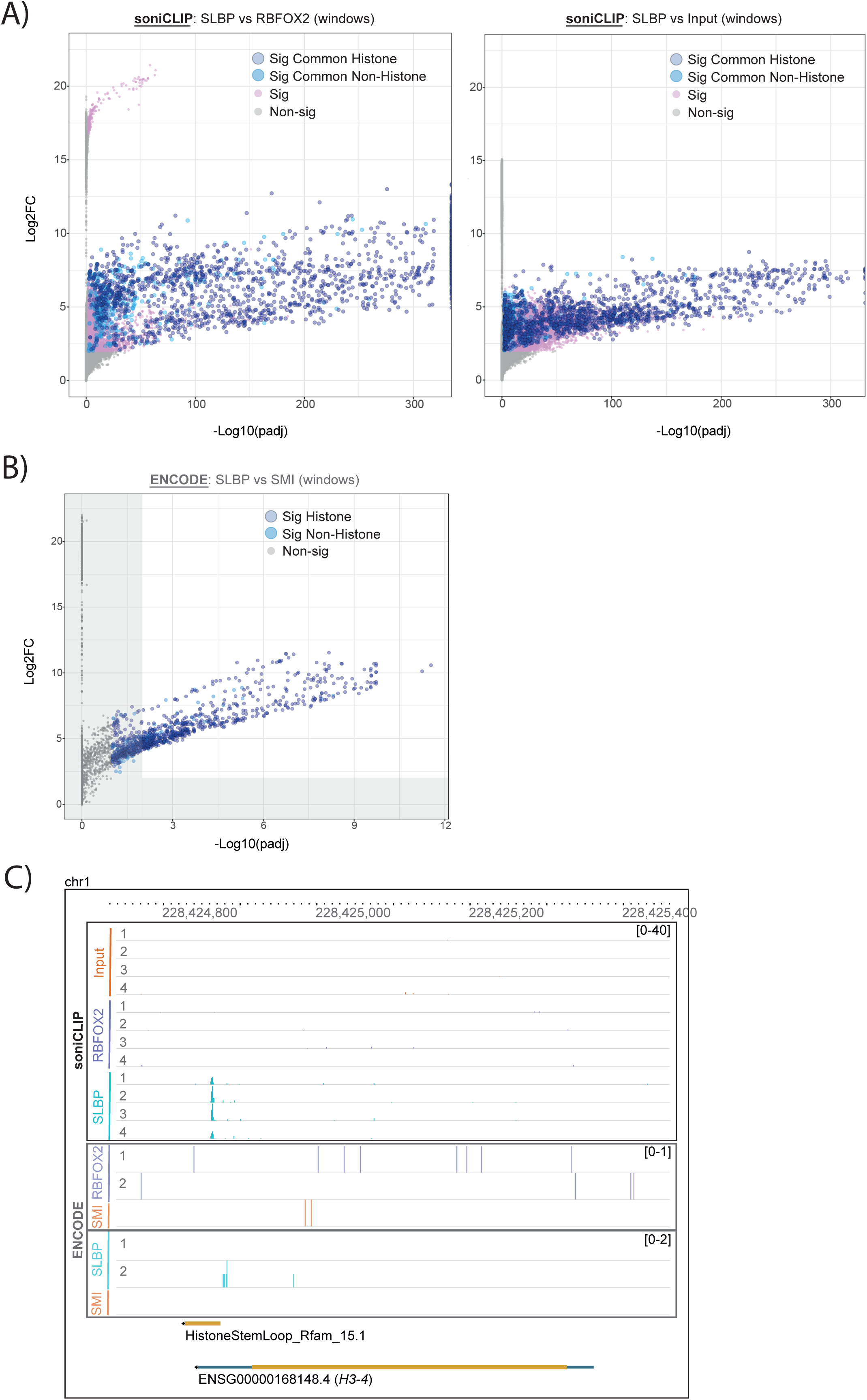
Identification of SLBP targets by soniCLIP recovers SLBP-bound histone stem-loop targets from low-input material. (A) Windows in soniCLIP analysis. Significant windows are highlighted (padj ≤0.01, L2FC ≥2), respectively, and the common histone and non-histone windows are depicted (overlap of SLBP IP vs RBFOX2 IP and SLBP IP vs Input). SLBP vs RBFOX2 as well as SLBP vs Input is shown. (B) Windows in ENCODE eCLIP analysis. Significant windows are highlighted (grey scale: padj ≤0.1, L2FC ≥1; padj ≤0.01, L2FC ≥2), respectively, and the significant histone and non-histone windows are depicted (SLBP IP vs SMI). (E) Exemplary genome browser tracks showing projected crosslink sites for soniCLIP and ENCODE eCLIP data. *H3-4* is shown.

**Supplementary Figure 4:**
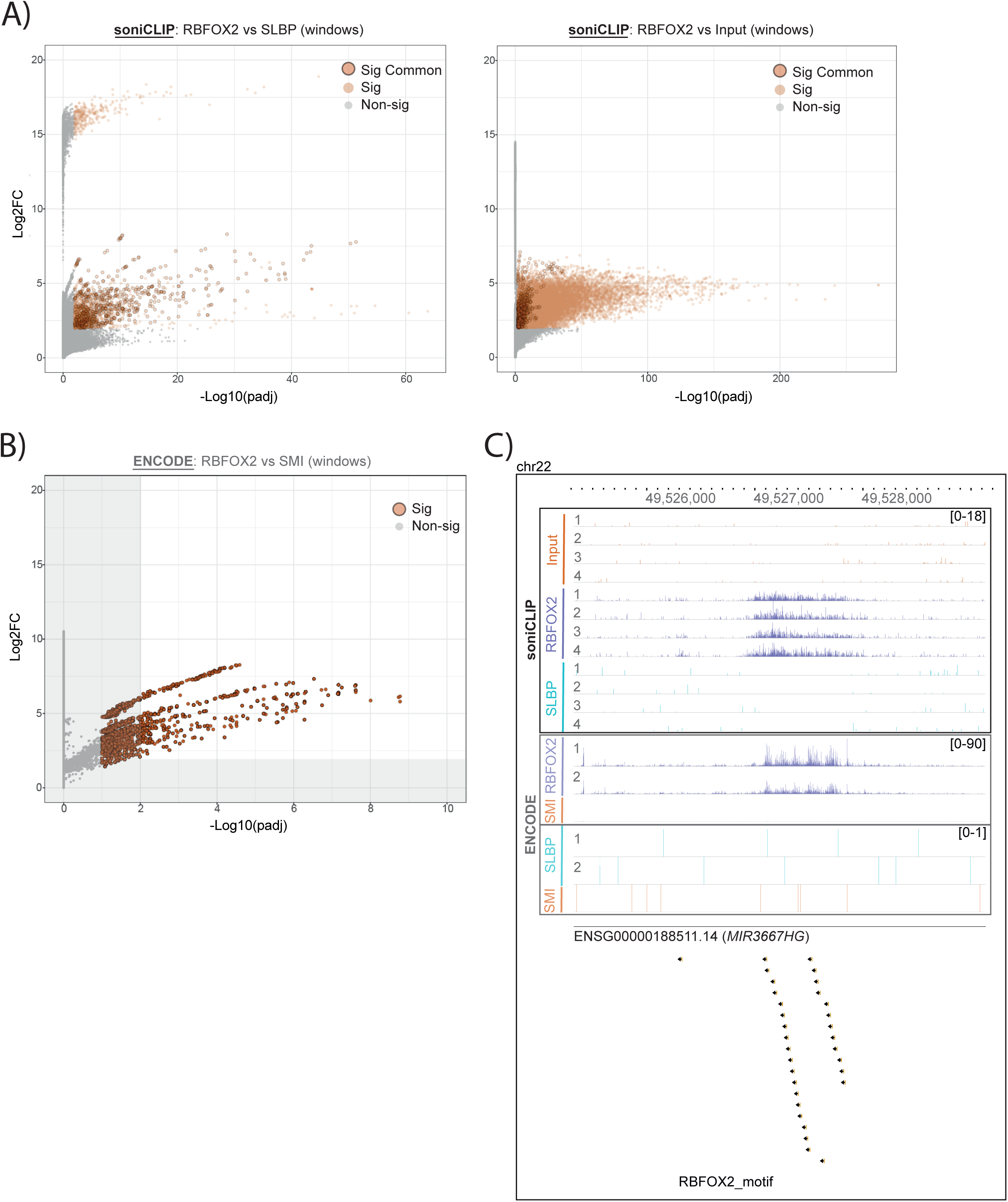
Identification of RBFOX2 motif-containing RNA regions with high specificity by soniCLIP from low-input material. (A) Windows in soniCLIP analysis. Significant windows are highlighted (padj ≤0.01, L2FC ≥2), respectively, and the common significant windows are depicted (overlap of RBFOX2 IP vs SLBP IP and RBFOX2 IP vs Input). RBFOX2 vs SLBP as well as RBFOX2 vs Input is shown. (B) Windows in ENCODE eCLIP analysis. Significant windows are highlighted (grey scale: padj ≤0.1, L2FC ≥1; padj ≤0.01, L2FC ≥2), respectively. RBFOX2 IP vs SMI is shown. (C) Exemplary genome browser tracks showing projected crosslink sites for soniCLIP and ENCODE eCLIP data. *MIR3667HG* is shown.

**Supplementary Figure 5:**
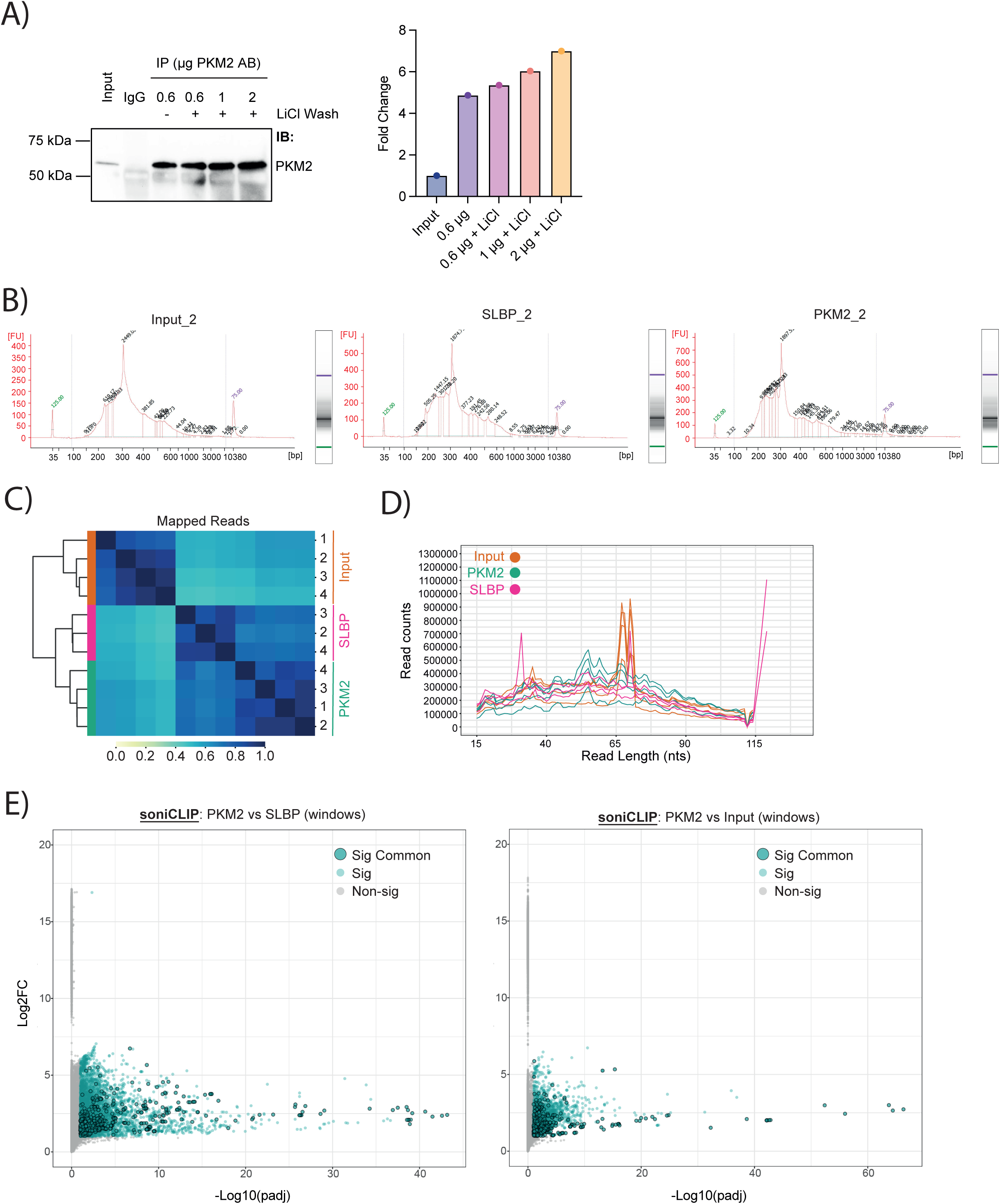
Robust target identification for the non-canonical RBP PKM2 by soniCLIP. (A) Immunoblot of immunoprecipitated PKM2 from HeLa cell lysates. N=1. (B) DNA Bioanalyzer profiles of soniCLIP libraries at the pre-size selection stage. Representative libraries are shown. N=4. (C) Sourmash plots of soniCLIP PKM2-SLBP dataset genome mapped reads as a means to visualize sequence similarity. Hierarchical clustering of similarity profiles scale 0-1. (D) Read length distribution soniCLIP PKM2-SLBP dataset genome mapped reads. (E) Windows in soniCLIP analysis. Significant windows are highlighted (padj ≤0.1, L2FC ≥1), respectively, and the common significant windows are depicted (overlap of PKM2 IP vs SLBP IP and PKM2 IP vs Input). PKM2 vs SLBP as well as RBPKM2 FOX2 vs Input is shown.

## SUPPLEMENTAL MATERIAL

### Buffers

Protease Inhibitor: cOmplete™, EDTA-free Protease Inhibitor Cocktail (Merck)

| Buffer | Composition |
| --- | --- |
| 2xD buffer | 200 mM KCl |
|  | 40 mM HEPES |
|  | 0.4 mM EDTA |
|  | 40% Glycerol |
|  | in deionized water; pH = 7.5 |
| 5x PNK buffer (pH 6.5) | 350 mM Tris-HCl |
|  | 50 mM MgCl<sub>2</sub> |
|  | in deionized water; pH = 6.5 |
| Dialysis buffer | 50 mM HEPES |
|  | 300 mM NaCl |
|  | 10% Glycerol |
|  | in deionized water, pH = 7.4 |
| EMSA binding reaction buffer | 4 mg/mL Ultrapure BSA (Merck #126609) |
|  | 1 mM DTT |
|  | 25 mM MgCl<sub>2</sub> |
|  | 20 µg/ml RNase Inhibitor |
|  | 100 mM KCl |
|  | 20 mM HEPES |
|  | 0.2 mM EDTA |
|  | 20% Glycerol |
|  | in deionized water |
| EMSA protein buffer | 25 mM HEPES |
|  | 150 mM KCl |
|  | 10% Glycerol |
|  | 5 mM MgCl<sub>2</sub> |
|  | 1 mM DTT |
|  | in deionized water; pH = 7.4 |
| Equilibration buffer | 20 mM Tris-HCl |
|  | 300 mM NaCl |
|  | 20 mM imidazole |
|  | 10% glycerol |
|  | In deionized water, pH = 8.0 |
| FastAP buffer | 100 mM KCl |
|  | 10 mM Tris-HCl |
|  | 0.02% Triton X-100 |
|  | 5 mM MgCl<sub>2</sub> |
|  | in deionized water |
| High-salt wash buffer | 500 mM NaCl |
|  | 50 mM Tris-HCl |
|  | 0.1% SDS |
|  | 0.5% Sodium Deoxycholate |
|  | 1% NP40 |
|  | 1X Protease Inhibitor |
|  | in deionized water |
| High-salt wash buffer LiCl | 80 mM LiCl |
|  | 500 mM NaCl |
|  | 50 mM Tris-HCl |
|  | 0.1% SDS |
|  | 0.5% Sodium Deoxycholate |
|  | 1% NP40 |
|  | 1X Protease Inhibitor |
|  | in deionized water |
| iCLIP100 lysis buffer | 100 mM NaCl |
|  | 50 mM Tris-HCl |
|  | 0.1% SDS |
|  | 0.5% Sodium Deoxycholate |
|  | 1% NP40 |
|  | 1X Protease Inhibitor |
|  | 55 µg/ml RNase inhibitor |
|  | in deionized water |
| Ion exchange equilibration buffer | 50 mM HEPES |
|  | 50 mM NaCl |
|  | 10% glycerol |
|  | In deionized water, pH = 7.4 |
| Native gel running buffer (10x) | 0.34 M Tris-HCl |
|  | 0.66 M HEPES |
|  | 1 mM EDTA |
|  | in deionized water; pH = 7.5 |
| PNK high-salt buffer | 1 M NaCl |
|  | 50 mM Tris-HCl |
|  | 0.1% SDS |
|  | 0.5% Sodium Deoxycholate |
|  | 1% NP40 |
|  | 1 mM EDTA |
|  | 1X Protease Inhibitor |
|  | in deionized water |
| PNK lysis buffer | 20 mM Tris pH 7.5 |
|  | 150 mM NaCl |
|  | 1% Triton |
|  | 1X Protease Inhibitor |
|  | 55 µg/ml RNase inhibitor |
|  | in deionized water |
| PNK wash buffer | 50 mM NaCl |
|  | 50 mM Tris-HCl |
|  | 10 mM MgCL2 |
|  | 0.5% NP40 |
|  | 1 mM EDTA |
|  | 1X Protease Inhibitor |
|  | in deionized water |
| Protein production lysis buffer | 50 mM Tris-HCl |
|  | 750 mM NaCl |
|  | 20 mM Imidazole |
|  | 10% Glycerol |
|  | 1X Protease Inhibitor |
|  | in deionized water, pH = 8.0 |
| Proteinase-K buffer qPCR | 30 mM Tris-HCl |
|  | 1% SDS |
|  | 10 mM EDTA |
|  | in deionized water |
| Proteinase-K buffer soniCLIP | 100 mM Tris-HCl |
|  | 50 mM NaCl |
|  | 10 mM EDTA |
|  | 0.2% SDS |
|  | in deionized water |
| RIP-qPCR wash buffer | 10 mM Tris-HCl |
|  | 0.2% Tween®20 |
|  | 1 mM EDTA |
|  | 1X Protease Inhibitor |
|  | in deionized water |
| RIPA | 150 mM NaCl |
|  | 50 mM Tris-HCl |
|  | 0.1% SDS |
|  | 0.5% Sodium Deoxycholate |
|  | 1% NP40 |
|  | 1 mM MgCl<sub>2</sub> |
|  | 0.1 mM CaCl<sub>2</sub> |
|  | 1X Protease Inhibitor |
|  | in deionized water |
| RIPA high-salt | 500 mM NaCl |
|  | 50 mM Tris-HCl |
|  | 0.1% SDS |
|  | 0.5% Sodium Deoxycholate |
|  | 1% NP40 |
|  | 1X Protease Inhibitor |
|  | in deionized water |
| RIPA LiCl high-salt | 80 mM LiCl |
|  | 500 mM NaCl |
|  | 50 mM Tris-HCl |
|  | 0.1% SDS |
|  | 0.5% Sodium Deoxycholate |
|  | 1% NP40 |
|  | 1X Protease Inhibitor |
|  | in deionized water |
| RIPA qPCR lysis buffer | 150 mM NaCl |
|  | 50 mM Tris-HCl |
|  | 0.1% SDS |
|  | 0.5% Sodium Deoxycholate |
|  | 1% NP40 |
|  | 1X Protease Inhibitor |
|  | in deionized water |
| SEC equilibration buffer | 25 mM HEPES |
|  | 150 mM KCl |
|  | 5 mM MgCl<sub>2</sub> |
|  | 10% Glycerol |
|  | 1 mM DTT |
|  | in deionized water, pH = 7.4 |
| T4 RNA ligation buffer without DTT | 50 mM Tris-HCl |
|  | 10 mM MgCl<sub>2</sub> |
|  | pH 7.5@25°C |
| Wash buffer | 20 mM Tris-HCl |
|  | 0.2% Tween®20 |
|  | 10 mM MgCl<sub>2</sub> |
|  | 1X Protease Inhibitor in deionized water |

### RNA oligos

Cyanine 5 (Cy5)-labelled RNA-oligonucleotides were acquired as RNase-free high-performance liquid chromatography (HPLC) purity grade from Integrated DNA Technologies (IDT) or Sigma Aldrich. Non-targeting oligonucleotides were selected with respect to their GC-content from the respective UTR regions. Sequences are always indicated 5’ - 3’.

Cy5-*IER3*_3‘-UTR

CGUGAGAUCCUUCCAUCUUCUUGAAGUCGCCUUUAGGGUG

Cy5-*FASN*_3‘-UTR_GC-matched_*IER3* (Ctrl) UAUUUAUUGCAUUGCUGGUAGAGACCCCCAGGCCUGUCCA

Cy5-*IER3*_3‘-UTR_GC-matched_*IER3* (Ctrl) AUCCGAAAAACCACAAAGAAACACCAGGCGUACCUGGUGC

Cy5-*MRFAP1L1*_5‘-UTR AGGCUUACAAUUAAAAGGAAGAAAAAAAAAUAAAGAUAAUUCGGG

Cy5-*FASN*_3‘-UTR_GC-matched_ *MRFAP1L1* (Ctrl) UUUGUUUUUCAAGAAAUGAUUCAAAUUGCUGCUUGGAUUUUGAAA

Cy5-*FASN*_unrelated (Ctrl) CUGGACUCGCUCAUGAGCGUGGAGGUGCGCCAGAC

### Vector Maps

The human wild-type PKM2 clone (#OHu20299) in vector pET24a was custom-generated and acquired from Thermo fisher (GeneArt Synthesis).

pET24a_hPMK2 (ORF clone (OHu20299; NM_002654.6)) 5‘-

TGGCGAATGGGACGCGCCCTGTAGCGGCGCATTAAGCGCGGCGGGTGTGGTGGTTACGCGCAGCGTGACCGCTACACTTGCCAGCGCCCTAGCGCCCGCTCCTTTCGCTTTCTTCCCTTCCTTTCTCGCCACGTTCGCCGGCTTTCCCCGTCAAGCTCTAAATCGGGGGCTCCCTTTAGGGTTCCGATTTAGTGCTTTACGGCACCTCGACCCCAAAAAACTTGATTAGGGTGATGGTTCACGTAGTGGGCCATCGCCCTGATAGACGGTTTTTCGCCCTTTGACGTTGGAGTCCACGTTCTTTAATAGTGGACTCTTGTTC CAAACTGGAACAACACTCAACCCTATCTCGGTCTATTCTTTTGATTTATAAGGGATTTTGCCGATTTCGGCCTATTGGTTAAAAAATGAGCTGATTTAACAAAAATTTAACGCGAATTTTAACAAAATATTAACGTTTACAATTTCAGGTGGCACTTTTCGGGGAAATGTGCGCGGAACCCCTATTTGTTTATTTTTCTAAATACATTCAAATATGTATCCGCTCATGAATTAATTCTTAGAAAAACTCATCGAGCATCAAATGAAACTGCAATTTATTCATATCAGGATTATCAATACCATATTTTTGAAAAAGCCGTTTCTGTAATGAAGGAGAAAACTCACCGAGGCAGTTCCATAGGATGGCAAGATCCTGGTATCGGTCTGCGATTCCGACTCGTCCAACATCAATACAACCTATTAATTTCCCCTCGTCAAAAATAAGGTTATCAAGTGAGAAATCACCATGAGTGACGACTGAATCCGGTGAGAATGGCAAAAGTTTATGCATTTCTTTCCAGACTTGTTCAACAGGCCAGCCATTACGCTCGTCATCAAAATCACTCGCATCAACCAAACCGTTATTCATTCGTGATTGCGCCTGAGCGAGACGAAATACGCGATCGCTGTTAAAAGGACAATTACAAACAGGAATCGAATGCAACCGGCGCAGGAACACTGCCAGCGCATCAACAATATTTTCACCTGAATCAGGATATTCTTCTAATACCTGGAATGCTGTTTTCCCGGGGATCGCAGTGGTGAGTAACCATGCATCATCAGGAGTACGGATAAAATGCTTGATGGTCGGAAGAGGCATAAATTCCGTCAGCCAGTTTAGTCTGACCATCTCATCTGTAACATCATTGGCAACGCTACCTTTGCCATGTTTCAGAAACAACTCTGGCGCATCGGGCTTCCCATACAATCGATAGATTGTCGCACCTGATTGCCCGACATTATCGCGAGCCCATTTATACCCATATAAATCAGCATCCATGTTGGAATTTAATCGCGGCCTAGAGCAAGACGTTTCCCGTTGAATATGGCTCATAACACCCCTTGTATTACTGTTTATGTAAGCAGACAGTTTTATTGTTCATGACCAAAATCCCTTAACGTGAGTTTTCGTTCCACTGAGCGTCAGACCCCGTAGAAAAGATCAAAGGATCTTCTTGAGATCCTTTTTTTCTGCGCGTAATCTGCTGCTTGCAAACAAAAAAACCACCGCTACCAGCGGTGGTTTGTTTGCCGGATCAAGAGCTACCAACTCTTTTTCCGAAGGTAACTGGCTTCAGCAGAGCGCAGATACCAAATACTGTCCTTCTAGTGTAGCCGTAGTTAGGCCACCACTTCAAGAACTCTGTAGCACCGCCTACATACCTCGCTCTGCTAATCCTGTTACCAGTGGCTGCTGCCAGTGGCGATAAGTCGTGTCTTACCGGGTTGGACTCAAGACGATAGTTACCGGATAAGGCGCAGCGGTCGGGCTGAACGGGGGGTTCGTGCACACAGCCCAGCTTGGAGCGAACGACCTACACCGAACTGAGATACCTACAGCGTGAGCTATGAGAAAGCGCCACGCTTCCCGAAGGGAGAAAGGCGGACAGGTATCCGGTAAGCGGCAGGGTCGGAACAGGAGAGCGCACGAGGGAGCTTCCAGGGGGAAACGCCTGGTATCTTTATAGTCCTGTCGGGTTTCGCCACCTCTGACTTGAGCGTCGATTTTTGTGATGCTCGTCAGGGGGGCGGAGCCTATGGAAAAACGCCAGCAACGCGGCCTTTTTACGGTTCCTGGCCTTTTGCTGGCCTTTTGCTCACATGTTCTTTCCTGCGTTATCCCCTGATT CTGTGGATAACCGTATTACCGCCTTTGAGTGAGCTGATACCGCTCGCCGCAGCCGAACGACCGAGCGCAGCGAGTCAGTGAGCGAGGAAGCGGAAGAGCGCCTGATGCGGTATTTTCTCCTTACGCATCTGTGCGGTATTTCACACCGCATATATGGTGCACTCTCAGTACAATCTGCTCTGATGCCGCATAGTTAAGCCAGTATACACTCCGCTATCGCTACGTGACTGGGTCATGGCTGCGCCCCGACACCCGCCAACACCCGCTGACGCGCCCTGACGGGCTTGTCTGCTCCCGGCATCCGCTTACAGACAAGCTGTGACCGTCTCCGGGAGCTGCATGTGTCAGAGGTTTTCACCGTCATCACCGAAACGCGCGAGGCAGCTGCGGTAAAGCTCATCAGCGTGGTCGTGAAGCGATTCACAGATGTCTGCCTGTTCATCCGCGTCCAGCTCGTTGAGTTTCTCCAGAAGCGTTAATGTCTGGCTTCTGATAAAGCGGGCCATGTTAAGGGCGGTTTTTTCCTGTTTGGTCACTGATGCCTCCGTGTAAGGGGGATTTCTGTTCATGGGGGTAATGATACCGATGAAACGAGAGAGGATGCTCACGATACGGGTTACTGATGATGAACATGCCCGGTTACTGGAACGTTGTGAGGGTAAACAACTGGCGGTATGGATGCGGCGGGACCAGAGAAAAATCACTCAGGGTCAATGCCAGCGCTTCGTTAATACAGATGTAGGTGTTCCACAGGGTAGCCAGCAGCATCCTGCGATGCAGATCCGGAACATAATGGTGCAGGGCGCTGACTTCCGCGTTTCCAGACTTTACGAAACACGGAAACCGAAGACCATTCATGTTGTTGCTCAGGTCGCAGACGTTTTGCAGCAGCAGTCGCTTCACGTTCGCTCGCGTATCGGTGATTCATTCTGCTAACCAGTAAGGCAACCCCGCCAGCCTAGCCGGGTCCTCAACGACAGGAGCACGATCATGCGCACCCGTGGGGCCGCCATGCCGGCGATAATGGCCTGCTTCTCGCCGAAACGTTTGGTGGCGGGACCAGTGACGAAGGCTTGAGCGAGGGCGTGCAAGATTCCGAATACCGCAAGCGACAGGCCGATCATCGTCGCGCTCCAGCGAAAGCGGTCCTCGCCGAAAATGACCCAGAGCGCTGCCGGCACCTGTCCTACGAGTTGCATGATAAAGAAGACAGTCATAAGTGCGGCGACGATAGTCATGCCCCGCGCCCACCGGAAGGAGCTGACTGGGTTGAAGGCTCTCAAGGGCATCGGTCGAGATCCCGGTGCCTAATGAGTGAGCTAACTTACATTAATTGCGTTGCGCTCACTGCCCGCTTTCCAGTCGGGAAACCTGTCGTGCCAGCTGCATTAATGAATCGGCCAACGCGCGGGGAGAGGCGGTTTGCGTATTGGGCGCCAGGGTGGTTTTTCTTTTCACCAGTGAGACGGGCAACAGCTGATTGCCCTTCACCGCCTGGCCCTGAGAGAGTTGCAGCAAGCGGTCCACGCTGGTTTGCCCCAGCAGGCGAAAATCCTGTTTGATGGTGGTTAACGGCGGGATATAACATGAGCTGTCTTCGGTATCGTCGTATCCCACTACCGAGATATCCGCACCAACGCGCAGCCCGGACTCGGTAATGGCGCGCATTGCGCCCAGCGCCATCTGATCGTTGGCAACCAGCATCGCAGTGGGAACGATGCCCTCATTCAGCATTTGCATGGTTTGTTGAAAACCGGACATGGCACTCCAGTCGCCTTCCCGTTCCGCTATCGGCTGAATTTGATTGCGAGTGAG ATATTTATGCCAGCCAGCCAGACGCAGACGCGCCGAGACAGAACTTAATGGGCCCGCTAACAGCGCGATTTGCTGGTGACCCAATGCGACCAGATGCTCCACGCCCAGTCGCGTACCGTCTTCATGGGAGAAAATAATACTGTTGATGGGTGTCTGGTCAGAGACATCAAGAAATAACGCCGGAACATTAGTGCAGGCAGCTTCCACAGCAATGGCATCCTGGTCATCCAGCGGATAGTTAATGATCAGCCCACTGACGCGTTGCGCGAGAAGATTGTGCACCGCCGCTTTACAGGCTTCGACGCCGCTTCGTTCTACCATCGACACCACCACGCTGGCACCCAGTTGATCGGCGCGAGATTTAATCGCCGCGACAATTTGCGACGGCGCGTGCAGGGCCAGACTGGAGGTGGCAACGCCAATCAGCAACGACTGTTTGCCCGCCAGTTGTTGTGCCACGCGGTTGGGAATGTAATTCAGCTCCGCCATCGCCGCTTCCACTTTTTCCCGCGTTTTCGCAGAAACGTGGCTGGCCTGGTTCACCACGCGGGAAACGGTCTGATAAGAGACACCGGCATACTCTGCGACATCGTATAACGTTACTGGTTTCACATTCACCACCCTGAATTGACTCTCTTCCGGGCGCTATCATGCCATACCGCGAAAGGTTTTGCGCCATTCGATGGTGTCCGGGATCTCGACGCTCTCCCTTATGCGACTCCTGCATTAGGAAGCAGCCCAGTAGTAGGTTGAGGCCGTTGAGCACCGCCGCCGCAAGGAATGGTGCATGCAAGGAGATGGCGCCCAACAGTCCCCCGGCCACGGGGCCTGCCACCATACCCACGCCGAAACAAGCGCTCATGAGCCCGAAGTGGCGAGCCCGATCTTCCCCATCGGTGATGTCGGCGATATAGGCGCCAGCAACCGCACCTGTGGCGCCGGTGATGCCGGCCACGATGCGTCCGGCGTAGAGGATCGAGATCTCGATCCCGCGAAATTAATACGACTCACTATAGGGGAATTGTGAGCGGATAACAATTCCCCTCTAGAAATAATTTTGTTTAACTTTAAGAAGGAGATATACATATGGCGCATCACCATCACCATCACGATTACGATATCCCGACAACCGAAAACCTGTATTTTCAGGGCATGTCGAAGCCCCATAGTGAAGCCGGGACTGCCTTCATTCAGACCCAGCAGCTGCACGCAGCCATGGCTGACACATTCCTGGAGCACATGTGCCGCCTGGACATTGATTCACCACCCATCACAGCCCGGAACACTGGCATCATCTGTACCATTGGCCCAGCTTCCCGATCAGTGGAGACGTTGAAGGAGATGATTAAGTCTGGAATGAATGTGGCTCGTCTGAACTTCTCTCATGGAACTCATGAGTACCATGCGGAGACCATCAAGAATGTGCGCACAGCCACGGAAAGCTTTGCTTCTGACCCCATCCTCTACCGGCCCGTTGCTGTGGCTCTAGACACTAAAGGACCTGAGATCCGAACTGGGCTCATCAAGGGCAGCGGCACTGCAGAGGTGGAGCTGAAGAAGGGAGCCACTCTCAAAATCACGCTGGATAACGCCTACATGGAAAAGTGTGACGAGAACATCCTGTGGCTGGACTACAAGAACATCTGCAAGGTGGTGGAAGTGGGCAGCAAGATCTACGTGGATGATGGGCTTATTTCTCTCCAGGTGAAGCAGAAAGGTGCCGACTTCCTGGTGACGGAGGTGGAAAATGGTGGCTCCTTGGGCAGCAAGAAGGGTGTGAACCTTCCTGGGGCTGCTGTGGACTTGCCTGCTGTGTC GGAGAAGGACATCCAGGATCTGAAGTTTGGGGTCGAGCAGGATGTTGATATGGTGTTTGCGTCATTCATCCGCAAGGCATCTGATGTCCATGAAGTTAGGAAGGTCCTGGGAGAGAAGGGAAAGAACATCAAGATTATCAGCAAAATCGAGAATCATGAGGGGGTTCGGAGGTTTGATGAAATCCTGGAGGCCAGTGATGGGATCATGGTGGCTCGTGGTGATCTAGGCATTGAGATTCCTGCAGAGAAGGTCTTCCTTGCTCAGAAGATGATGATTGGACGGTGCAACCGAGCTGGGAAGCCTGTCATCTGTGCTACTCAGATGCTGGAGAGCATGATCAAGAAGCCCCGCCCCACTCGGGCTGAAGGCAGTGATGTGGCCAATGCAGTCCTGGATGGAGCCGACTGCATCATGCTGTCTGGAGAAACAGCCAAAGGGGACTATCCTCTGGAGGCTGTGCGCATGCAGCACCTGATTGCCCGTGAGGCAGAGGCTGCCATCTACCACTTGCAATTATTTGAGGAACTCCGCCGCCTGGCGCCCATTACCAGCGACCCCACAGAAGCCACCGCCGTGGGTGCCGTGGAGGCCTCCTTCAAGTGCTGCAGTGGGGCCATAATCGTCCTCACCAAGTCTGGCAGGTCTGCTCACCAGGTGGCCAGATACCGCCCACGTGCCCCCATCATTGCTGTGACCCGGAATCCCCAGACAGCTCGTCAGGCCCACCTGTACCGTGGCATCTTCCCTGTGCTGTGCAAGGACCCAGTCCAGGAGGCCTGGGCTGAGGACGTGGACCTCCGGGTGAACTTTGCCATGAATGTTGGCAAGGCCCGAGGCTTCTTCAAGAAGGGAGATGTGGTCATTGTGCTGACCGGATGGCGCCCTGGCTCCGGCTTCACCAACACCATGCGTGTTGTTCCTGTGCCGTAATGAGTCGACAAGCTTGCGGCCGCACTCGAGCACCACCACCACCACCACTGAGATCCGGCTGCTAACAAAGCCCGAAAGGAAGCTGAGTTGGCTGCTGCCACCGCTGAGCAATAACTAGCATAACCCCTTGGGGCCTCTAAACGGGTCTTGAGGGGTTTTTTGCTGAAAGGAGGAACTATATCCGGAT-3’

